# A pharmacological toolkit for human microglia identifies Topoisomerase I inhibitors as immunomodulators for Alzheimer’s disease

**DOI:** 10.1101/2024.02.06.579103

**Authors:** Verena Haage, John F. Tuddenham, Natacha Comandante-Lou, Alex Bautista, Anna Monzel, Rebecca Chiu, Masashi Fujita, Frankie G. Garcia, Prabesh Bhattarai, Ronak Patel, Alice Buonfiglioli, Juan Idiarte, Mathieu Herman, Alison Rinderspacher, Angeliki Mela, Wenting Zhao, Michael G. Argenziano, Julia L. Furnari, Matei A. Banu, Donald W. Landry, Jeffrey N. Bruce, Peter Canoll, Ya Zhang, Tal Nuriel, Caghan Kizil, Andrew A. Sproul, Lotje D. de Witte, Peter A. Sims, Vilas Menon, Martin Picard, Philip L. De Jager

## Abstract

While efforts to identify microglial subtypes have recently accelerated, the relation of transcriptomically defined states to function has been largely limited to *in silico* annotations. Here, we characterize a set of pharmacological compounds that have been proposed to polarize human microglia towards two distinct states – one enriched for AD and MS genes and another characterized by increased expression of antigen presentation genes. Using different model systems including HMC3 cells, iPSC-derived microglia and cerebral organoids, we characterize the effect of these compounds in mimicking human microglial subtypes *in vitro*. We show that the Topoisomerase I inhibitor Camptothecin induces a CD74^high^/MHC^high^ microglial subtype which is specialized in amyloid beta phagocytosis. Camptothecin suppressed amyloid toxicity and restored microglia back to their homeostatic state in a zebrafish amyloid model. Our work provides avenues to recapitulate human microglial subtypes *in vitro*, enabling functional characterization and providing a foundation for modulating human microglia *in vivo*.

## Introduction

With the multiplicity of key functions carried out by microglia to maintain homeostasis in the central nervous system (CNS) ^1–3^, the characterization of specialized human microglial subsets with regards to brain region, function, physiological and pathophysiological context has been a subject of intense interest over the past decade ^4^. Improved single-cell (sc) RNA sequencing methods led to the recent emergence of several datasets characterizing human microglia and have transformed our understanding of human microglial heterogeneity ^4–12^.

While the currently available datasets propose different human microglial population structures, highlighting the limitations of current profiling technologies, certain patterns are emerging from these transcriptomic data, underscoring the need for a shared framework for nomenclature of human microglia. In particular, given the possible range of cellular metabolic states and mitochondrial phenotypes (i.e., mitotypes) that influence cellular behaviors ^13^, a frame of reference for profiling bioenergetic profiles among microglia is needed ^14^. The greater challenge however has been linking transcriptionally defined clusters of human microglia to key microglial functions. This is critical to understand both homeostatic functions and roles in the development and progression of neurodegenerative disease. Aligning transcriptional signatures with cellular function will also be crucial with respect to therapeutic development, to guide the modulation of selected microglial subtypes in different contexts.

Given the range of current human microglial subpopulation models, approaches to build *in vitro* models for specific human microglial subtypes in order to study their function in depth have been limited ^12,15^. Here, we deeply characterize our prior *in silico* analysis-driven pharmacologic approach of recapitulating certain human microglia subtypes in model systems to enable their functional evaluation. Specifically, the current report started at the end of our prior effort (Tuddenham et al., 2022) which (1) proposed a cross-disease human microglial atlas comprised of 12 subtypes, (2) prioritized 14 compounds for microglial polarization after an *in silico* drug screening approach, and (3) identified 3 candidate compounds which mimicked selected human microglial subtypes in the Human Microglial Cell 3 (HMC3) cell line *in vitro*.

In this study, we further characterized the 3 compounds Narciclasine, Torin2 and Camptothecin as foundational compounds of a pharmacological toolbox with which to study the function of human microglial subpopulations *in vitro* and *in vivo*. Through structure-activity relationship analysis and a series of scRNA sequencing experiments and functional assays using compound-treated microglial model systems including the HMC3 microglial cell line, iPSC-derived microglia (iMG), cerebral organoids with incorporated iMGs, and mice. we identified Camptothecin and its Food and Drug Agency (FDA)-approved analog Topotecan as potent drivers of an immunologically activated microglial subtype, characterized by high expression of CD74 and MHC I as well as MHC II (CD74^high^/MHC^high^ cells); these genes are enriched in antigen presenting cells (APC), including myeloid cells specialized for this function. These topoisomerase I inhibitors also suppress an SRGAP2^high^/MEF2A^high^ microglial subtype that is enriched in susceptibility genes for AD and MS (Disease Enriched Microglia, DEM)^12^. Treatment of an amyloidosis zebrafish model with Camptothecin reduced activated microglia and prevented reduced synaptic density in zebrafish treated amyloid b, a key proteoform for the amyloid proteinopathy that, with taupathy, defines Alzheimer’s disease (AD).

## Results

### Pharmacologic engagement of two distinct transcriptomic signatures in HMC3 microglia

Given that Camptothecin reduced the DEM signature and enhanced an APC signature, it was of great interest from a therapeutic development perspective: a compound with this property could theoretically be useful in shifting the distribution of human microglial subtypes *in vivo* away from disease-enriched SRGAP2^high^/MEF2A^high^ microglia and towards the CD74^high^/MHC^high^ microglial subtype. On the other hand the other two compounds from our prior evaluation ^12^, Narciclasine and Torin 2 engaged different aspects of the SRGAP2^high^/MEF2A^high^ DEM signature, therefore inducing gene expression changes that were opposite to those of Camptothecin. The ability to polarize microglia towards the DEM subtype could be very useful to explore, mechanistically, how this subtype enriched for disease genes may be contributing to pathophysiology (**Figure 1A**).

**Figure 1.**
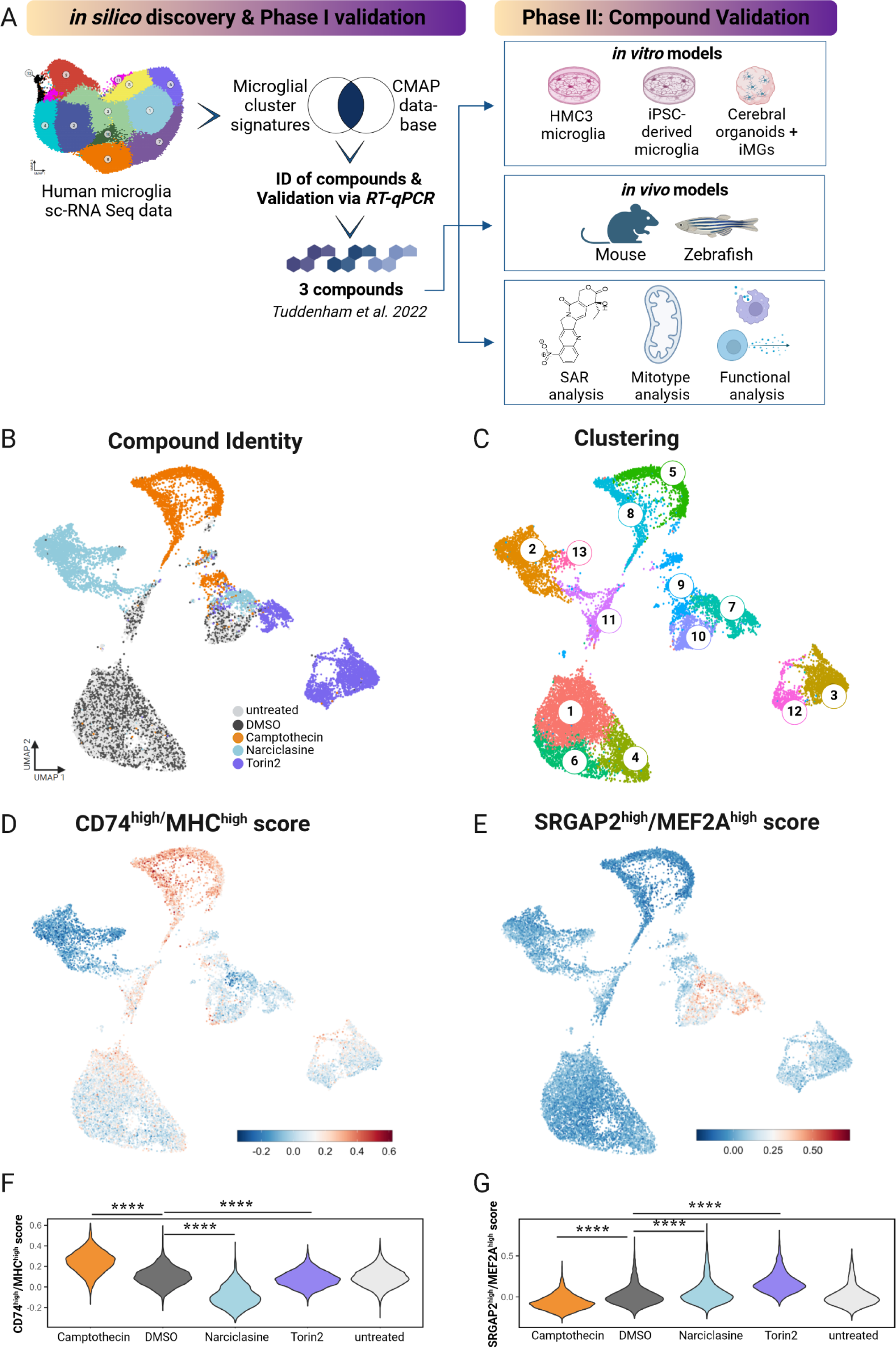
A.Overview of the study design and sc-RNA sequencing analysis of compound-treated HMC3 microglia. Schematic diagram depicting *in silico* discovery phase, **phase I** of compound validation via RT-qPCR expression analysis, **phase II** of compound validation for three selected compounds Narciclasine, Torin2, Camptothecin in various *in vitro* microglial model systems and *in vivo* models. In **validation phase III** structure activity relationship analysis (SAR), mitochondrial phenotyping (mitotype) and functional assays were performed. **B. UMAP of compound-treated HMC3s showing treatment.** Each dot is a single cell, colored by compound treatment condition - untreated (light grey), DMSO control (dark grey), Camptothecin (orange) Narciclasine (light blue), Torin2 (purple). **C. UMAP of compound-treated HMC3s showing cluster identity.** Each dot is a single cell, colored by cluster identity ranging from clusters 1-13. **D. CD74^high^/MHC^high^ signature in Camptothecin- treated HMC3 microglia.** Enrichment of the top 50 cluster genes was calculated on a per-cell basis compared to background genes with similar expression levels. Cells are colored by log- fold change of the CD74^high^/MHC^high^ gene set. **E. SRGAP2^high^/MEF2A^high^ signature in compound-treated HMC3 microglia. F.** Violin plots depict the per-cell **CD74^high^/MHC^high^ module score** grouped by drug treatment. For statistical analysis pairwise Wilcoxon rank-sum tests were performed, ****p.adj ≤ 0.0001. **G.** Violin plots show **SRGAP2^high^/MEF2A^high^ module score**/cell grouped by drug treatment. For statistical analysis pairwise Wilcoxon sum-rank tests were performed, ****p.adj ≤ 0.0001.

### Single cell level characterization of the three tool compounds

To determine the extent of heterogeneity among HMC3 cells at baseline and after exposure to our compounds, we repeated our previously established treatment paradigm^12^ by exposing HMC3 microglia to Narciclasine (0.1µM), Torin2 (10µM), Camptothecin (1µM), or DMSO, and we left some untreated as well. After 24hrs, cells were labelled with cell hashing antibodies, pooled and scRNAseq was performed (**Figure 1B**). Each compound polarized HMC3 cells into a distinct transcriptional space (**Figure 1B**) and distinct clusters were identified in association with different treatment conditions (**Figure 1C**).

To assess whether each compound induced the respective target transcriptional signature, we assessed the top 100 differentially expressed genes in each microglial subtype in treated HMC3 microglia and colored each cell by the level of expression for the CD74^high^/MHC^high^ (**Figure 1D**) or the SRGAP2^high^/MEF2A^high^ (**Figure 1E**) signatures. Camptothecin clearly induced the CD74^high^/MHC^high^ signature, and Camptothecin-treated cells were noticeably depleted in the SRGAP2^high^/MEF2A^high^ signature (**Figure 1D-E**). Torin2 and Narciclasine induce a modest enrichment of the SRGAP2^high^/MEF2A^high^ signature in the main set of cells derived from each of these two conditions; however the SRGAP2^high^/MEF2A^high^ signature is much more strongly induced in the small subset of Torin2- and Narciclasine-treated cells found in the center of the projection which likely stem from a transcriptionally discrete minor subtype of untreated and DMSO-treated cells (**Figure 1B**), suggesting that this subtype of HMC3 cells may have a more permissive context for expression of the DEM (SRGAP2^high^/MEF2A^high^) signature. **Figure 1F-G** depicts a quantitative analysis of our data, showing that Camptothecin significantly increases the expression of the CD74^high^/MHC^high^ signature; Torin 2 and Narciclasine both significantly reduce this signature. The inverse is seen for SRGAP2^high^/MEF2A^high^ signature. These results are consistent with our predictions.

### Structure Activity Relationship (SAR) Analysis identifies Topotecan as a Camptothecin- analog in inducing the CD74^high^/MHC^high^ microglial signature

Next, we performed structure activity relationship (SAR) analysis by selecting three to four analog compounds with either a similar chemical structure or the same target protein for each of our three compounds of interest: Narciclasine, Torin2 and Camptothecin (**Figure 2A**).

**Figure 2.**
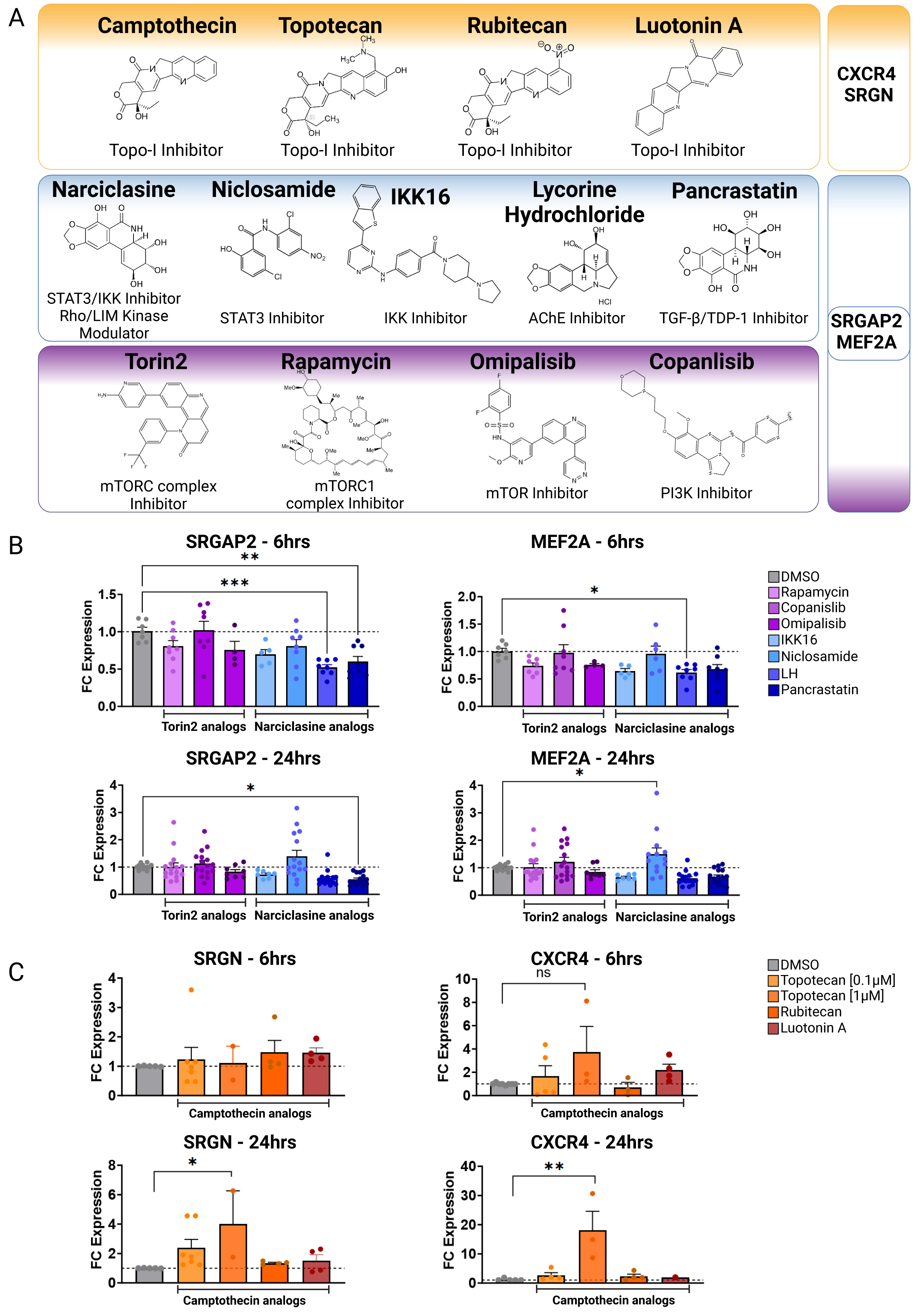
Structure activity relationship (SAR) analysis. **A.** Overview of chemical structures and functional targets of selected compounds for structure activity relationship analysis. B. Marker gene expression (*SRGN, CXCR4*) in HMC3 microglia treated with Camptothecin analogs (6hrs and 24hrs) assessed via RT-qPCR. CT values were normalized to *HPRT1*. Bars represent fold change expression (mean ± SEM) in relation to DMSO control. For statistical analysis, one-way ANOVA followed by Dunnett’s multiple comparisons test was performed. *p.adj ≤ 0.05; **p.adj ≤ 0.01; ***p.adj ≤ 0.001; ****p.adj ≤ 0.0001. **C. Marker gene expression (*SRGAP2*, *MEF2A*) in HMC3 microglia treated with analog compounds for Torin2 and Narciclasine (6hrs and 24hrs),** assessed via RT-qPCR. CT values were normalized to housekeeping gene *HPRT1*. Bars represent fold change expression (mean ± SEM) in relation to DMSO control. For statistical analysis, one-way ANOVA followed by Dunnett’s multiple comparisons test was performed. *p.adj ≤ 0.05; **p.adj ≤ 0.01; ***p.adj ≤ 0.001; ****p.adj ≤ 0.0001.

For the LIM/ROCK and STAT3/NFKB pathway inhibitor Narciclasine, we selected Lycorine Hydrochloride and Pancrastatin as structural analogs, as well as the IKK inhibitors IKK16 and Niclosamide as functional analogs. For Torin2, we selected the original mTORC1 inhibitor Rapamycin as a functional analog, Copanlisib, a PI3 Kinase inhibitor, as a structural analog and Omipalisib as a functional and structural analog. For Camptothecin, we selected three different structural and functional analogs: Rubitecan, Topotecan, and Luotonin A (**Figure 2A**).

Following titration experiments to determine the optimal dose for each compound (**Figure S2**), HMC3 microglia were treated with the selected doses (6hrs, 24hrs) and marker expression was assessed via RT-qPCR followed by calculation of the log fold change (FC) expression of each marker gene in comparison to DMSO-treated control (**Figure 2B-C**). *SRGAP2* and *MEF2A* expression was assessed to measure the SRGAP2^high^/MEF2A^high^ microglial signature, while *CXCR4* and *SRGN* were used as markers to determine the CD74^high^/MHC^high^ microglial signature.

<u>Narciclasine:</u> While Lycorine hydrochloride unexpectedly downregulated *MEF2A* and *SRGAP2* significantly after 6hrs, this effect was transient and not present at 24 hours (**Figure 2B**). Pancrastatin, showed a consistent downregulation of *SGRAP2* 6hrs and 24hrs following treatment of HMC3 microglia, which however was not observed for the second marker gene *MEF2A*. The only compound that induced an upregulation of one of the marker genes, *MEF2A*, following 24hrs after treatment is Niclosamide, a functional analog of Narciclasine. These data suggest that the observed effect of Narciclasine in inducing the SRGAP2^high^/MEF2A^high^ signature in HMC3 microglia, may depend on its STAT3 and NFKB inhibitor function rather than on its chemical structure.

Torin2: none of the selected compounds - Rapamycin, Copanlisib and Omipalisib - produced the expected changes in *MEF2A* and *SRGAP2* expression (**Figure 2B**) ^12^.

<u>Camptothecin</u>: we identified Topotecan as a clear driver of the CD74^high^/MHC^high^ signature (**Figure 2C**; *CXCR4* and *SRGN* expression was upregulated 24hrs following treatment), in particular with the higher dosage of 1µM tested. These results of Topotecan inducing the CD74^high^/MHC^high^ microglial subtype are of particular interest as Topotecan is a more effective and safer version of Camptothecin that is FDA approved for the treatment of small-cell lung cancer ^16^.

Overall, from the structure activity relationship analysis, we identified Niclosamide as a potentially interesting second compound mimicking the effects of Narciclasine to induce the SRGAP2^high^/MEF2A^high^ human microglial subcluster ^12^ *in vitro* and Topotecan as a potent driver of the CD74^high^/MHC^high^ signature.

Studies exploring the role of topoisomerase I in neuroinflammation have increased over the past few years ^17^, and we generated an additional dataset suggesting an interesting role for Topotecan with regards to the CD74^high^/MHC^high^ subtype. We assessed scRNAseq data generated from of a series of experiments performed on primary human GBM surgical excisions treated exogenously with Topotecan for 18 hours ^18,19^. Myeloid cells were then sorted *in silico,* and analysis of Topotecan response was performed at the level of individual samples and the aggregate of all Topotecan-treated samples (**Figure S2D**). When performing GSEA, in two out of the three examined human-derived glioblastoma brain slice samples treated with Topotecan, we observed an upregulation of the human microglial CD74^high^/MHC^high^ signature as well as a downregulation of the SRGAP2^high^/MEF2A^high^ signature. These data align with our HMC3 observations, and the data generated from this complex model systems provide strong support for the idea that this effect of Topotecan signal is robust, reproducible, and present in primary human cells derived from neurosurgical specimen.

### Compound-treated iMG and CO-iMG confirm Camptothecin as a potent tool compound to induce the CD74^high^/MHC^high^ signature

Next, we tested our compounds in induced-pluripotent stem cell (iPSC)-derived microglia-like cells (iMG) by exposing iMG to Narciclasine, Torin2, Camptothecin or its analog Topotecan (**Figure 2C**) in addition to the DMSO control on day 28-29 of differentiation^20^. After 24 hours of compound exposure, cells were collected, and scRNAseq data were generated (**Fig. 3A, Experiment 1**).

**Figure 3.**
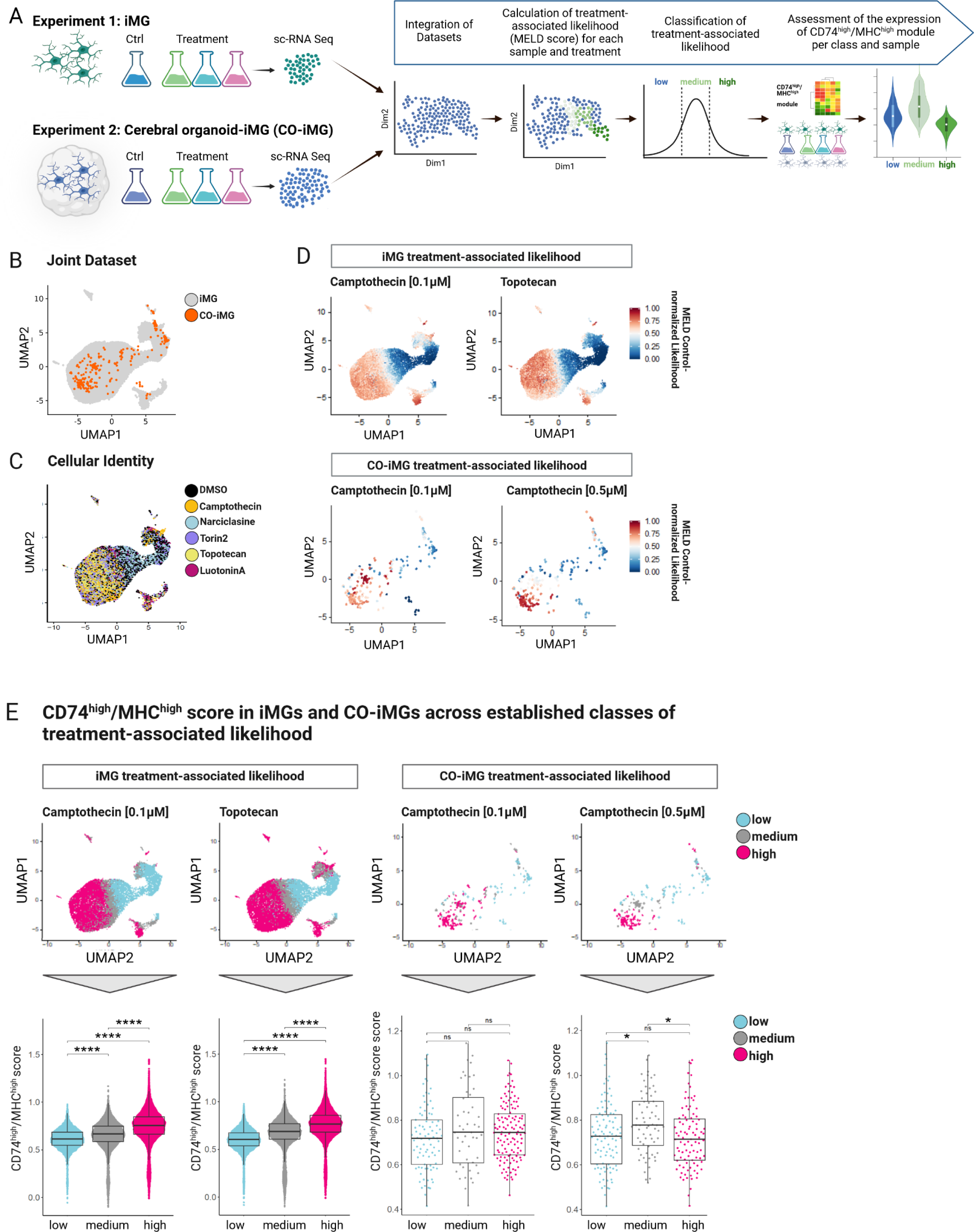
**sc-RNA sequencing of compound-treated iMG and CO-iMG reveals upregulation of the CD74^high^/MHC^high^ module following Camptothecin and Topotecan treatment. A**. **Overview of experimental design**. iPSC-derived microglia (iMG), treated with compounds (Narciclasine, Torin2, Camptothecin, Topotecan, LuotoninA; Experiment 1) and cerebral organoids containing implanted iMG (CO-iMG), treated with compounds (Camptothecin 0.1µM and 0.5µM) or DMSO as control (24hrs) were subjected to sc-RNA sequencing. CO-iMG were projected onto the iMG data and MELD ^23^ was used to quantify the relative treatment-associated likelihood for individual cells. Cells were then classified based on the relative treatment-associated likelihood (low; medium; high) and expression of CD74^high^/MHC^high^ signature genes was assessed across the three levels of relative treatment-associated likelihood calculated for each drug. **B**. UMAP of the integrated datasets (Seurat query-mapping pipeline) of compound-treated iMGs (gray) and CO-iMGs (orange). Each dot represents one cell. **C**. UMAP of integrated datasets (Seurat query-mapping pipeline) derived from controls and compound-treated iMGs and CO-iMGs. Each dot represents a cell from different treatment conditions before MELD score analysis (DMSO; Torin2; Narciclasine; Camptothecin; Topotecan; LuotoninA). **D. Individual iMGs colored by treatment-associated likelihood in the joint UMAP.** Same UMAP as in B-C, except that cells are colored by the relative treatment-associated likelihood scores (red:high, blue:low) calculated by MELD for treatment conditions: Camptothecin- and Topotecan treatment in iMGs, and Camptothecin treatment at 0.1µM and 0.5µM in CO-iMGs. **E. CD74^high^/MHC^high^ score in iMGs and CO-iMGs across different levels of treatment-associated likelihood.** Top row - same UMAP as in B-C, with cells colored by three different levels (low, medium, high) of treatment-associated likelihood for each drug-treatment condition. Bottom row - single-cell distribution of the CD74^high^/MHC^high^ module scores across the three levels of treatment-associated likelihood for each of the given samples. Each dot represents a single cell, statistical analysis was performed using unpaired t-test; ns = non-significant; ****p ≤ 0.0001. Each boxplot highlights the median, lower and upper quartiles. Whiskers indicate 1.5 times interquartile ranges.

Along with these experiments, we performed a second line of experiments in a more complex model system: cerebral organoids co-cultured with isogenic iMG (CO-iMG), an advanced model systems in which microglia can be studied in a human 3 dimensional microenvironment (see **Methods**)^21^. In short, CO were first differentiated from iPSC (25-35 days old), and hematopoietic progenitor cells (HPC, 12 days old) differentiated from iPSC were then seeded onto the CO. The CO and HPC were then co-cultured for 7 days, allowing the HPC to differentiate into microglia (iMG) and integrate into the cerebral organoids (CO-iMG). On day 7 after co-culture, CO-iMG were treated with Narciclasine (0.01µM), Torin2 (10nM), Camptothecin (0.1µM, 0.5µM) or DMSO for 24hrs (for dose-titration on iMG, see **Fig. S3A**) followed by processing for scRNAseq (**Fig. 3A, Experiment 2**). As for Narciclasine and Torin2 treatment, we did not recover sufficient CO-iMG cells for analysis and therefore focused on a joint analysis of the datasets derived from Camptothecin- and Topotecan-treated iMG and Camptothecin-treated CO-iMG (for Narciclasine- and Camptothecin-treated iMG data see **Figure S3C**. First, we computationally isolated CO-iMG cells for further analysis based on the co-expression pattern of canonical microglial markers (see **Methods**) in individual cells (**Figure 3A**, Experiment 2). We then interrogated the effect of the selected compounds on the transcriptional states of iMG and CO-iMG to assess the extent to which the expected drug effects are recapitulated in these model systems. To answer these questions, we first projected both datasets onto an integrated space in an unsupervised manner ^22^ using the iMG data as reference and the CO-iMG data as query, such that cells from both systems could be jointly analyzed. **Figure 3B** shows the Uniform Manifold Approximation and Projection (UMAP) embeddings of the iMG and the CO-iMG projected onto the same space. Most CO-iMG were able to be mapped within the iMG space, suggesting transcriptional similarity between the two model systems.

To quantify the effect of compound-treatment across conditions and model systems at the single-cell level, we applied the MELD algorithm ^23^ on the integrated iMG and CO-iMG dataset. MELD captures the continuous and heterogeneous nature of cell populations in response to perturbations, and it models the treatment-associated relative likelihood for each cell from a given condition compared to the control. **Figure 3C** shows the treatment-associated likelihood for individual cells in the iMG and CO-iMG integrated space.

While the treatment of iMG with Narciclasine barely induced transcriptional changes in the iMG, Torin2-treatment showed definitive changes as indicated by the cells with a high control-normalized treatment-associated likelihood for this compound (**Figure S3C-D**). These data, however, were not further followed up on as Torin2-treatment was toxic to CO-iMG. Camptothecin-treatment of iMG and CO-iMG yielded the most interesting results. Based on the calculated treatment-associated likelihood of iMG associated with 0.1µM Camptothecin treatment, cells with the highest likelihood to be derived from the treatment condition clustered mostly on the left part of the UMAP (**Figure 3D)**. Similarly, CO-iMG projected onto the same UMAP spaces showed similar distributions of Camptothecin-associated likelihood for both doses, with a continuous increase from the right to left side of the UMAP. In short, our data derived from Camptothecin-treated CO-iMG resemble closely those of Camptothecin-treated iMG, underlining the robustness of Camptothecin in inducing a specific transcriptional program across different human microglial model systems. We subsequently defined three classes (low, medium, high) of treatment-associated likelihood for each experiment and treatment condition by fitting the data with a three-class gaussian mixture model (**Figure 3E**, see **Methods**) and assessed the expression of the CD74^high^/MHC^high^ signature. While 0.1µM Camptothecin-treated iMG significantly increased CD74^high^/MHC^high^ module expression correlating with the increase of treatment-associated likelihood from low to medium to high classes. The CO-iMG data are quite sparse, and while we see a trend for higher CD74^high^/MHC^high^ signature in the cells of the higher likelihood of treatment effect classes, it was only significant in the medium likelihood classes with 0.5µM Camptothecin treatment compared to the low likelihood cell class (**Figure S3E**). Deeper sequencing of CO-iMG or a greater number of treated CO-iMG will be needed to fully demonstrate the effect of Camptothecin on the CD74^high^/MHC^high^ signature in this model system so that enough iMG can be recovered.

In iMG treated with Topotecan, we observed a highly similar pattern in the distribution of cells as for Camptothecin-treated iMG (**Figure 3D**). Strikingly, Topotecan induced the CD74^high^/MHC^high^ signature as strongly as Camptothecin, further highlighting Topotecan as an interesting, FDA-approved drug mimicking the observed effects of Camptothecin in polarizing microglia-like cells (**Figure 3E**).

### Functional characterization: *in vitro* polarized cells differ in their endocytic and phagocytic properties

One of the key functions of microglia involves clearance of cellular debris, pathogen-infected cells or complement-tagged synapses from live neurons^24^ to maintain brain homeostasis. Here, we chose to use HMC3 cells and a series of substrates to assess different phagocytic phenotypes: (1) pHrodo Dextran as a readout for endocytosis and macropinocytosis, (2) fluorescently-labeled Aβ 1-42 as a readout for phagocytosis in an AD context, and (3) pHrodo labeled heat-inactivated E.coli as a readout for Toll-like-receptor-mediated phagocytosis in an inflammatory context ^25^ (**Figure 4A**). Cytochalasin D, an inhibitor of actin polymerization, was used as a negative control (**Figure S4A**). Interestingly, we observed distinct endocytic and phagocytic phenotypes depending on the state of HMC3 polarization (**Figure 4B-D**).

**Figure 4.**
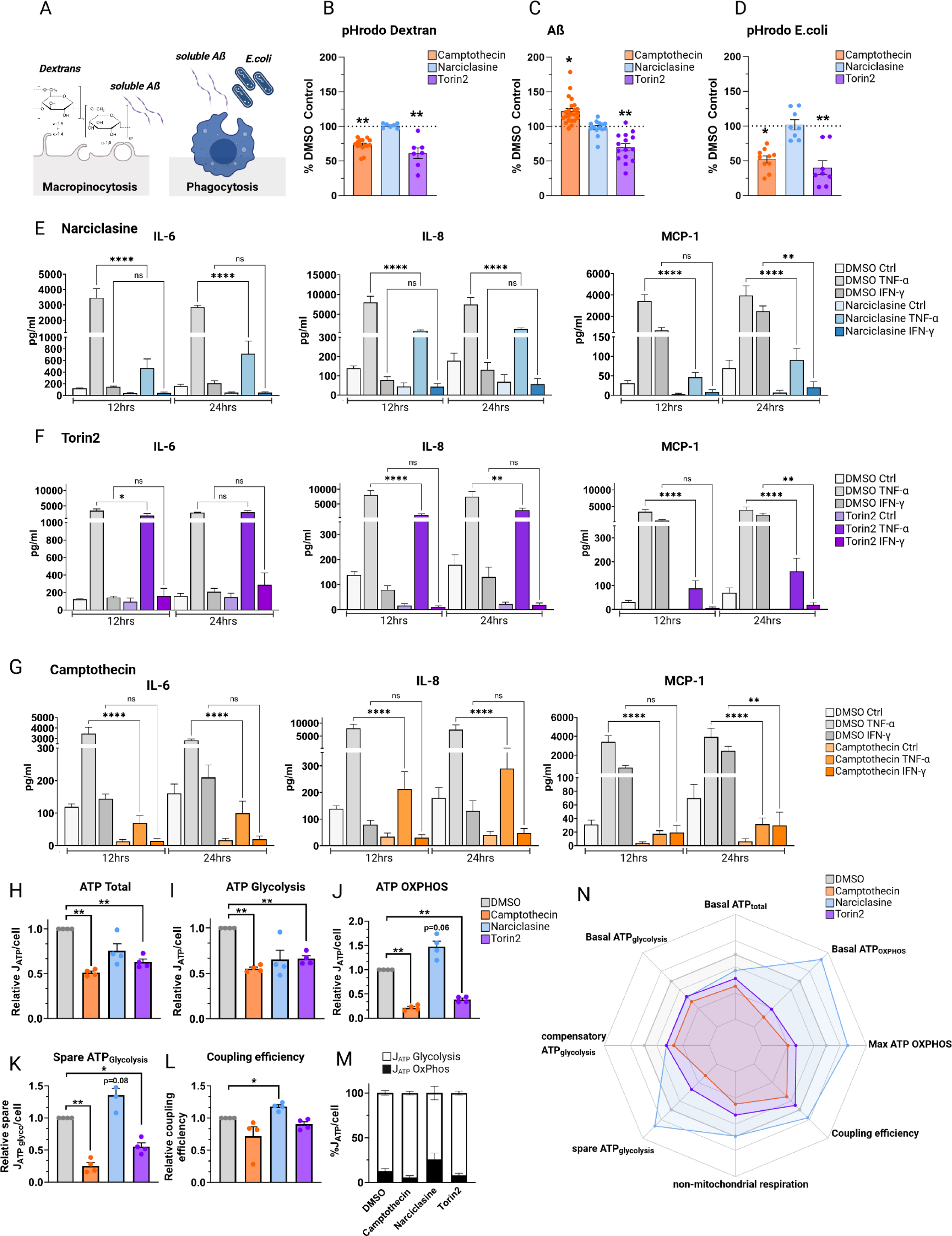
Compound-treated HMC3 microglia exhibit substrate-specific endocytic and phagocytic phenotypes and differences in secretion of pro-inflammatory cytokines. **A-C. Phagocytic phenotypes. A.** Graph depicting nature of the different assays assessing macropinocytosis (pHrodo Dextran, soluble Aß) and phagocytosis (Aß, E.coli). B. Vorinostat and Entinostat upregulate pHrodoDextran phagocytosis. HMC3 microglia, pretreated with respective compounds or DMSO as control (24hrs), were exposed to pHrodo-labeled Dextran (**B**), AlexaFluor 647-labeled Aβ (**C**) or pHrodo-labelled E.coli particles (**D**) for 1hr, uptake was assessed via flow cytometry. Each dot represents one independent experiment (mean ± SEM; Camptothecin – orange; Narciclasine – blue; Torin2 - purple). Phagocytosis was normalized to percent DMSO control, for statistical analysis, log-fold change values in comparison to DMSO-treated control samples were analyzed using one-way ANOVA followed by Dunnett’s multiple comparison test. *p.adj ≤ 0.05; **p.adj ≤ 0.01. **E.-G. Cytokine secretion of compound- or DMSO-pretreated HMC3 cells** (24hrs) followed by stimulation with either TNF-a (0.3 µg/mL), IFN-y (0.3 µg/mL) or H2O as control for 12 or 24hrs. Pro-inflammatory cytokine secretion was assessed using a human pro-inflammatory cytokine discovery assay. Bar graphs depict measured amount of cytokines IL-6, IL-8, MCP-1 (mean ± SEM) in pg/ml for DMSO control-treated samples (white, light grey, grey) or compound-treated samples (light blue, purple, orange). For statistical analysis, one-way ANOVA followed by Tukey’s multiple comparisons test with a single pooled variance was performed. *p.adj ≤ 0.05; **p.adj ≤ 0.01; ***p.adj ≤ 0.001; ****p.adj ≤ 0.0001. **H-N**. **MitoStress test on HMC3 cells treated with Camptothecin, Narciclasine, and Torin2,** depicting ATP total/cell (**H**), ATP Glycolysis (**I**), ATP generated through OxPhos (**J**), Spare ATP Glycolysis (**K**), coupling efficiency (**L**) and ATP generated by Glycolysis or OxPhos relative to DMSO (**M**). **N**. Data show means ± SEM. *p.adj ≤ 0.05, **p.adj ≤ 0.01, ***p.adj ≤ 0.001 test with BH adjustment. HedgesG for effect size.

Narciclasine, did not induce any changes in the assessed phagocytic phenotypes in comparison to DMSO-treated controls (**Figure 4B**), while Torin2 treatment - causing mTOR inhibition - induced a downregulation in uptake activity for all three substrates. Thus, Torin 2 affects endocytic as well as phagocytic processes (**Figure 4B-D**), in line with previous reports of impaired microglial uptake in mTOR knockout mice ^26^ as well as an increase in microglial- mediated ß-amyloid plaque clearance upon mTOR activation in the 5xFAD mouse model ^27^. These distinct effects on uptake by Narciclasine and Torin2 - despite driving overlapping transcriptomic and proteomic phenotypes in HMC3 microglia^12^ – are consistent with our scRNAseq data (**Figure 1**) which revealed that these compounds polarize HMC3 cells into largely distinct clusters.

On the other hand, Camptothecin (**Figure 4B-D**) led to a reduction in endocytic activity (lower Dextran uptake) as well as a downregulation of TLR-mediated phagocytosis of E.coli while upregulating uptake of monomeric Aβ. This pathway-specific effect of Camptothecin suggests a specialization of this human microglial *in vitro* cell model to a microglial subtype with a specific phagocytic function, one that may be very relevant in the context of Aβ clearance in incipient AD.

Thus, our uptake assays have provided a validation of our experimental strategy of using repurposed compounds to drive polarization of microglia-like cells towards pre-defined transcriptionally distinct subtype which reflect functionally distinct states. All three compounds have a clearly distinct uptake pattern for the selected substrates and help to prioritize Camptothecin as a lead compound for further development in the realm of AD.

### CD74^high^/MHC^high^ and SRGAP2^high^/MEF2A^high^ microglia show reduced secretion of pro-inflammatory chemokines and cytokines

Activating immune responses is another important function of microglia, so we evaluated the effect of each compound in modulating response to TNF-α or IFN-γ stimulation (**Figure 4E-G, Figure S4**) by measuring 15 pro-inflammatory cytokines in culture supernatants (see **Methods**). Across the three different compounds, changes were observed with Interleukin-6 (IL-6), Interleukin 8 (IL-8 or CXCL8), and the chemokine monocyte chemoattractant protein 1 (MCP-1 or CCL2), known to affect microglia ^28^.

Narciclasine- and Torin2-treated HMC3 cells showed similar results with regards to IL-8 and MCP-1 secretion when stimulated with TNF-α or IFN-γ, but they differed in the secretion of IL- 6 following TNF-α stimulation (**Figure 4E-G**). TNF-α-induced IL-8 and MCP-1 secretion was significantly reduced 12 and 24hrs after treatment with both Narciclasine and Torin2 treatment, suggesting an effect for both compounds on TNF-α signaling. MCP-1 levels were also reduced 24hrs after IFN-γ stimulation, suggesting that this pathway may also be affected by these two compounds. Narciclasine also strongly reduced IL-6 secretion following TNF-α stimulation (12hrs, 24hrs), whereas Torin2 showed a transient minor reduction in TNF-α-induced IL-6 secretion after 12hrs, but not 24hrs. Neither compound affected IFN-γ-induced IL-6 secretion (**Figure 4E-G**). Thus, the SRGAP2^high^/MEF2A^high^ subtype seems to primarily have an altered TNF-α response.

Camptothecin treatment strongly suppressed IL-6, IL-8 as well as MCP-1 secretion 12hrs and 24hrs following stimulation with TNF-α (**Figure 4G**). Moreover, Camptothecin reduced MCP-1 secretion downstream of IFN-γ signaling 24hrs following IFN-γ stimulation. These results are consistent with reports of Camptothecin (1) suppressing inflammatory gene expression including IL-8, IL-6 as well as MCP-1 secretion in different model systems ^17,29^, (2) inhibiting the activation of NFKB ^30^ and (3) polarizing microglia towards an anti-inflammatory phenotype^30^.

Interestingly, Topotecan treatment fully recapitulated the observed cytokine-release pattern induced by Camptothecin (**Figure 4G**), thereby functionally supporting the data generated as part of the SAR analysis and further positioning Topotecan as an interesting candidate for inducing the CD74^high^/MHC^high^ human microglial signature *in vitro* in future studies.

### Functional characterization: Mitochondrial phenotypes differ among *in vitro* polarized HMC3 microglia

Our recent report describing human microglial subpopulations also suggested a central metabolic divide between the two major poles of our human microglial population structure model ^12^, with microglial subtypes on the left side of the population structure (Cluster 4/9, **Figure 1A**) being enriched in genes related to oxidative phosphorylation (OxPhos) and the right side of the population structure corresponding to the SRGAP2^high^/MEF2A^high^ subtypes (Clusters 1, 6), being enriched for genes involved in alternative metabolic pathways and heterocyclic metabolism ^12^. Metabolic phenotypes have been previously linked to distinct microglial activation states with homeostatic microglia being most reliant on OxPhos; following activation, reactive microglia switch their metabolism to glycolysis, a faster but less efficient way to generate ATP ^31^.

We therefore assessed respiratory states of compound-treated HMC3 microglia with a MitoStressTest (**Methods**), 24hrs following treatment, and derived interpretable ATP production rates for OxPhos and glycolysis. All three compounds induced a hypometabolic state where the total ATP production rate (JATP-Total) relative to DMSO control was decreased by 25-49%. Camptothecin showed the strongest effect, followed by Torin2 and then Narciclasine (**Figure 4H**). Importantly, Camptothecin and Torin2 had an effect on both, JATP-Glycolysis and JATP-OxPhos, suggesting that these compounds induce a general hypometabolic state reflecting lower energy demand. Narciclasine treatment, however, induced a different metabolic phenotype. Even though total energy demand was reduced overall, the fraction of JATP-OxPhos relative to JATP-Glycolysis was shifted towards OxPhos, and JATP-OxPhos was even higher in Narciclasine treated cells than in DMSO control (p=0.06; **Figure 4H**). These data are in line with previous GO term analysis indicating transcriptional changes, specifically mitochondrial transcriptional changes, as well as changes in metabolism in Narciclasine-treated HMC3 microglia (**Figure 5C**).

**Figure 5.**
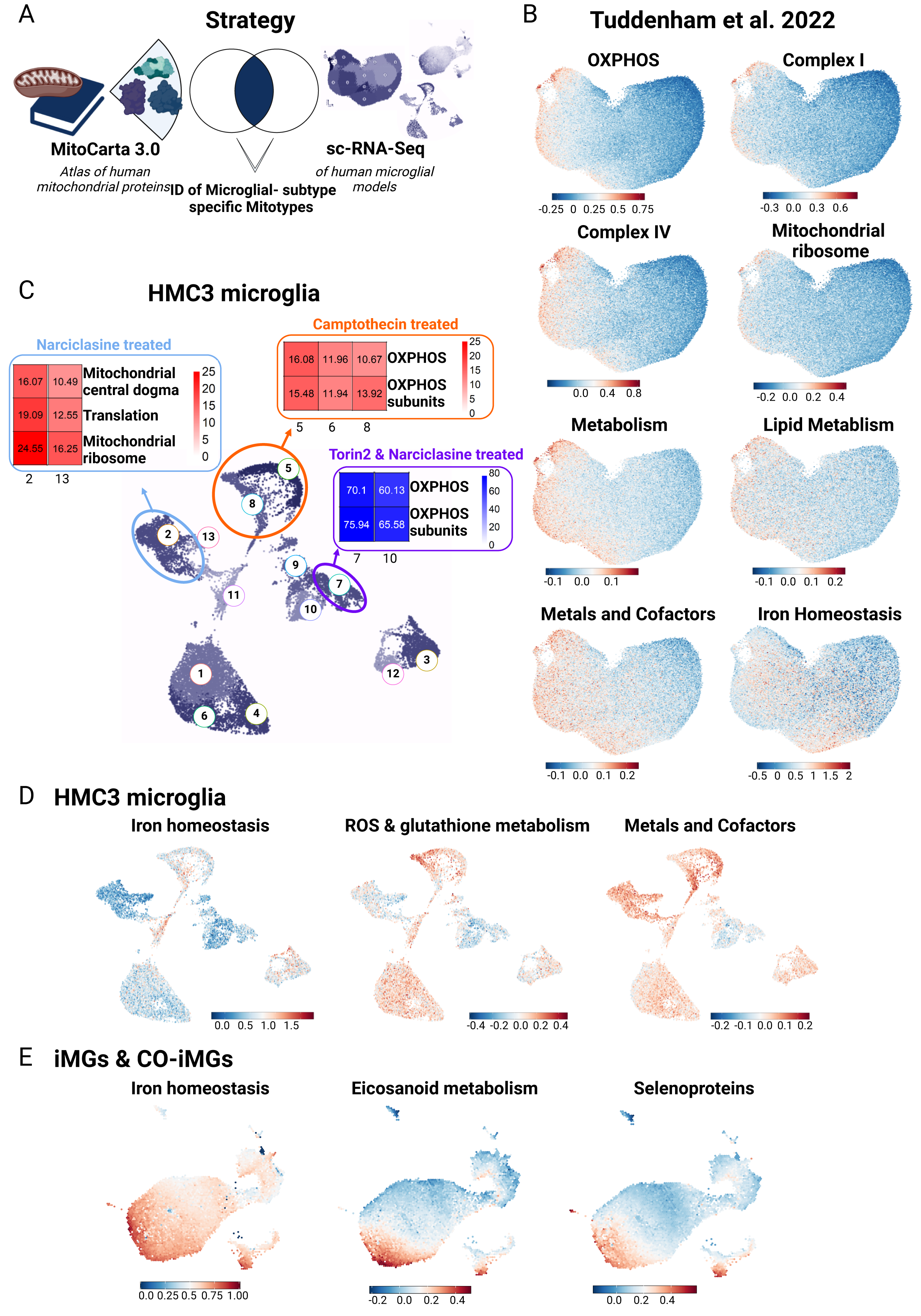
Mitochondrial phenotyping identifies association between distinct human microglial subtypes and specific mitochondrial phenotypes. A. Strategy for mitochondrial phenotyping analysis. 149 mitochondrial pathways defined from the MitoCarta 3.0 ^37^, were used to perform Gene Set Enrichment Analysis (GSEA) in the human microglial sc-RNA-Seq dataset ^12^ **B.** Selected results of GSEA for specific mitotypes in human microglial subtypes are shown. Legend depicts module score, showing average expression for module genes vs. a background of similarly expressed control genes. **C. Mitotype analysis results in compound-treated HMC3 microglia following GSEA for the 149 mitochondrial pathways from** ^37^. Genes from HMC3-treated sc-RNA Seq data were ordered based on average log-fold change between cells from the respective treatment condition vs. control. Enrichment of gene modules associated with different aspects of mitochondrial activity was performed in upregulated gene lists associated with each cluster using hypergeometric test with Benjamini-Hochberg correction. **D. Results of selected results of mitotype GSEA projected into UMAP of compound-treated HMC3 microglia.** Plots show most specific and distinct GSEA results for mitochondrial phenotypes in compound-treated HMC3 microglial clusters. Legend depicts module score, showing average expression for module genes versus a background of similarly expressed control genes. **E. Results of selected results of mitotype GSEA projected into joint UMAP of compound-treated iMG and CO-iMG.** Plots show most specific and distinct GSEA results for mitochondrial phenotypes in compound-treated iMG and CO-iMG microglial clusters. Legend depicts module score, showing average expression for module genes vs. a background of similarly expressed control genes.

When uncoupling mitochondrial respiration from ATP production, cells possess the ability to compensate for this lack of energy by increasing glycolysis. The difference between the compensatory glycolysis and baseline glycolysis levels is defined as the spare glycolytic capacity ^32^. Narciclasine treatment, which increased the OxPhos:Glycolysis ratio at baseline, also increases spare glycolytic capacity induced by ATP synthase inhibition. Interestingly, glycolytic capacity has been previously linked to cellular reprogramming ^33^ which is in line with our GO term analysis of Narciclasine-treated HMC3 cells (**Figure 5C**). Camptothecin and Torin2 treatment decreased the spare glycolytic capacity dramatically, suggesting that these cells have almost no scope to respond to an acute metabolic challenge. However, a glycolysis-specific stress test (glucose depletion and/or glycolysis inhibition) would be necessary to confirm this observation.

We further calculated the coupling efficiency, indicating the proportion of oxygen consumption used for ATP synthesis in comparison with the oxygen consumption rate (OCR) driving the proton leak (ATP-linked OCR/basal OCR)^34^, with proton leak defined as the remaining basal respiration not coupled to ATP production ^35^. While we observed an increase in the coupling efficiency in Narciclasine-treated HMC3 microglia, Camptothecin and Torin2 reduced mitochondrial coupling efficiency.

In sum, all three compounds altered the metabolic state of HMC3 in addition to altering this cell line’s transcriptome (**Figure 1**). While total ATP synthesis was reduced to a hypometabolic state upon treatment with these three compounds, Camptothecin- and Torin2- treated cells downregulated glycolysis, OxPhos and spare capacity while Narciclasine induced a metabolic switch from glycolysis towards OxPhos-dependent metabolism with an increase in coupling efficiency. These results thus further elaborate the functional nature of the distinct microglial states induced by polarization with our compounds.

### Human microglial subtypes exhibit enrichment of specific mitotypes

The distinct metabolic signatures in our microglia subtypes motivated us to examine whether these subtypes also showed distinct molecular mitochondrial signatures.^36^ We repurposed our scRNAseq data generated from living human microglia^12^ and used a computational approach of mitochondrial phenotyping (i.e. mitotyping) by deploying mitochondrial pathway annotations from the human MitoCarta3.0 resource, a catalogue comprised of 1136 human mitochondrial genes and 149 pathway annotations ^37^. We then calculated a Gene Set enrichment analysis (GSEA) scores to assess the enrichment of the 149 specific MitoPathways (**Table S2**) in the 12 human microglial subtypes that we previously defined^12^ (**Figure 1A**, **Figure 5A-B**) as well as in our compound-treated *in vitro* model systems (HMC3 microglia – **Figure 5D**; iMG & CO- iMG: **Figure 5E**).

Interestingly, we observed an enrichment of genes related to OxPhos on the left side of the human microglial population structure which contains homeostatic cells^12^ (**Figure 5B**), and, at the same time, OxPhos related MitoPathways including Complex I, III and IV were strongly downregulated on the right side of the population structure, consistent with prior observations from GO analyses of this dataset ^12^, suggesting that cells at opposite extremes of our diagram have distinct metabolic states. The right side of the diagram contains the SRGAP2^high^/MEF2A^high^ microglia (clusters 1 and 6).

In Narciclasine-treated HMC3, we noted earlier two induced subpopulations (**Figure 1B).** In the major subpopulation, we observed a strong enrichment of MitoPathways associated with mitochondrial central dogma, translation, mitochondrial ribosome and protein import and sorting, indicating changes in the mitochondrial DNA (**Figure 5C**), consistent with GSEA for the MitoCarta 3.0 gene sets in our previously published bulk RNA-Seq data of Narciclasine-treated HMC3 microglia (**Figure S5**)^12^. These data are in accordance with our observations from the seahorse analysis where we observed a metabolic switch from glycolysis to oxidative phosphorylation upon Narciclasine treatment in HMC3 microglia (**Figure 4I**). With regards to the previously published^12^ human microglial population structure, changes in the metabolic phenotype of microglia were expected when exposing the cells to Narciclasine, but we expected a shift away from OxPhos towards alternative heterocyclic pathways ^12^. As HMC3 microglia-like cells have a glycolytic phenotype at baseline conditions (**Figure 4I**) and are dividing cells, our results from the HMC3 model system may be skewed by the baseline state of these cells vs. primary microglia. Yet, what is consistent across both the model system and primary cells is that Narciclasine treatment induces a metabolic switch that seems to go different directions depending on the nature of the baseline state.

Interestingly, the second, minor subpopulation induced by Narciclasine (and also by Torin 2) (**Figure 1C**) treatment induced a small subset of cells which we consider to be the closest in mimicking the nature of the original human SRGAP2^high^/MEF2A^high^ microglial subset (**Figure 1B**). The mitochondrial phenotype of this specific cluster revealed downregulation of OxPhos and OxPhos subunits in comparison to control cells (clusters 1,6,4,10; **Figure 5C**) which is in line with the human dataset, where a decrease in OxPhos and an upregulation of alternative heterocyclic pathways was observed ^12^. Thus, this minor subpopulation induced by both Narciclasine and Torin 2 may be the best one to study going forwards; further work is needed to resolve whether these HMC3 cells were in a different baseline state – which may be the case given the distribution of cells - or simply represent a lower probability endpoint of differentiation following SRGAP2^high^/MEF2A^high^ signature induction, relative to the larger induced population which is metabolically (and transcriptionally) distinct.

On the other hand, the CD74^high^/MHC^high^ signature induced by Camptothecin, is not at the extreme ends of the major metabolic axis of our microglial population structure model (**Figure 1**); they appear to be a minor microglial subtype following a different vector of differentiation^12^. Using MitoPathway annotations, we detected an upregulation of OxPhos subunit and Complex I subunit pathways in the cells induced by Camptothecin (**Figure 5C**, clusters 5,8 in Figure 2B) in comparison to DMSO-treated control HMC3 microglia (**Figure 5C**, clusters 1,6,4,10 in Figure 2B). Repurposing the purified living primary human microglia data, the CD74^high^/MHC^high^ subtype (labeled as cluster 10 in the model in **Figure 1A**) displays enrichment of MitoPathways related to lipid metabolism, ROS and glutathione metabolism, as well as metals and cofactors. When performing the same type of analysis in our established compound-based models including HMC3 microglia, iMG and CO-iMG, we could recapitulate these findings in all our *in vitro* models where mitotype analysis of Camptothecin-treated HMC3 microglia as well as Camptothecin- or Topotecan-treated iMG and CO-iMG showed a higher degree of OxPhos and iron homeostasis (**Figure 5D-E**).

Overall, CD74^high^/MHC^high^ human microglia-like cells represent a state enriched for certain functions: (1) enhanced antigen presenting cell machinery, (2) preferential enhanced uptake of a subset of substrates (Ab), (3) diminished cytokine secretion response to TNF-α, and (4) enhanced OxPhos metabolism, reactive oxygen species (ROS) generation and glutathione metabolism as well as in metals and cofactors (**Figure 5D**), results confirming previous mitotype analysis in the primary microglia scRNAseq dataset (**Figure 5B**).

In sum, our mitochondrial phenotyping analysis in both, human microglial subpopulations as well as established compound-based *in vitro* models in HMC3 microglia suggest that distinct mitochondrial phenotypes are associated with specific, transcriptionally-defined microglial subpopulations.

### Narciclasine and Camptothecin induce transcriptional shifts in microglia *in vivo*

Next, we tested the efficacy of our pharmacological toolkit in an *in vivo* model system. We therefore treated wildtype C57BL/6 mice with two of our three compounds: Narciclasine polarizing microglia towards the SRGAP2^high^/MEF2A^high^ subtype and Camptothecin, for polarization towards the CD74^high^/MHC^high^ subtype. As both compounds have been previously been studied in the CNS in mice, we implemented prior treatment paradigms (**Methods**) ^30,38^. Narciclasine reduces pro-inflammatory cytokines and COX2 while inducing anti-inflammatory cytokines in the context of LPS stimulation ^38^. Camptothecin has been shown to exert neuroprotective effects in a mouse model of Parkinson’s disease (PD) and was proposed to polarize microglia towards an anti-inflammatory phenotype.^30^ Over the course of treatment with either drug, we did not observe any weight changes in female or male mice (**Figure S6A**).

At the end of each treatment paradigm, we sorted brain cell suspensions for CD11b^+^CD45^+^CX3CR1^+^ microglia and profiled them with scRNA-Seq (**Figure 6A, Fig S6B**). Microglia derived from controls and both drug treatments were analyzed jointly, but separately for each sex (**Figure 6A**). For each integrated dataset (female and male), we performed batch correction and calculated the treatment-associated likelihood using the MELD algorithm ^23^ (**Figure S6C**). Cells were subsequently grouped into classes ranging from low, medium to high treatment-associated likelihood, and the expression of both signatures, SRGAP2^high^/MEF2A^high^ and CD74^high^/MHC^high^ was assessed (**Figure 6C**).

**Figure 6.**
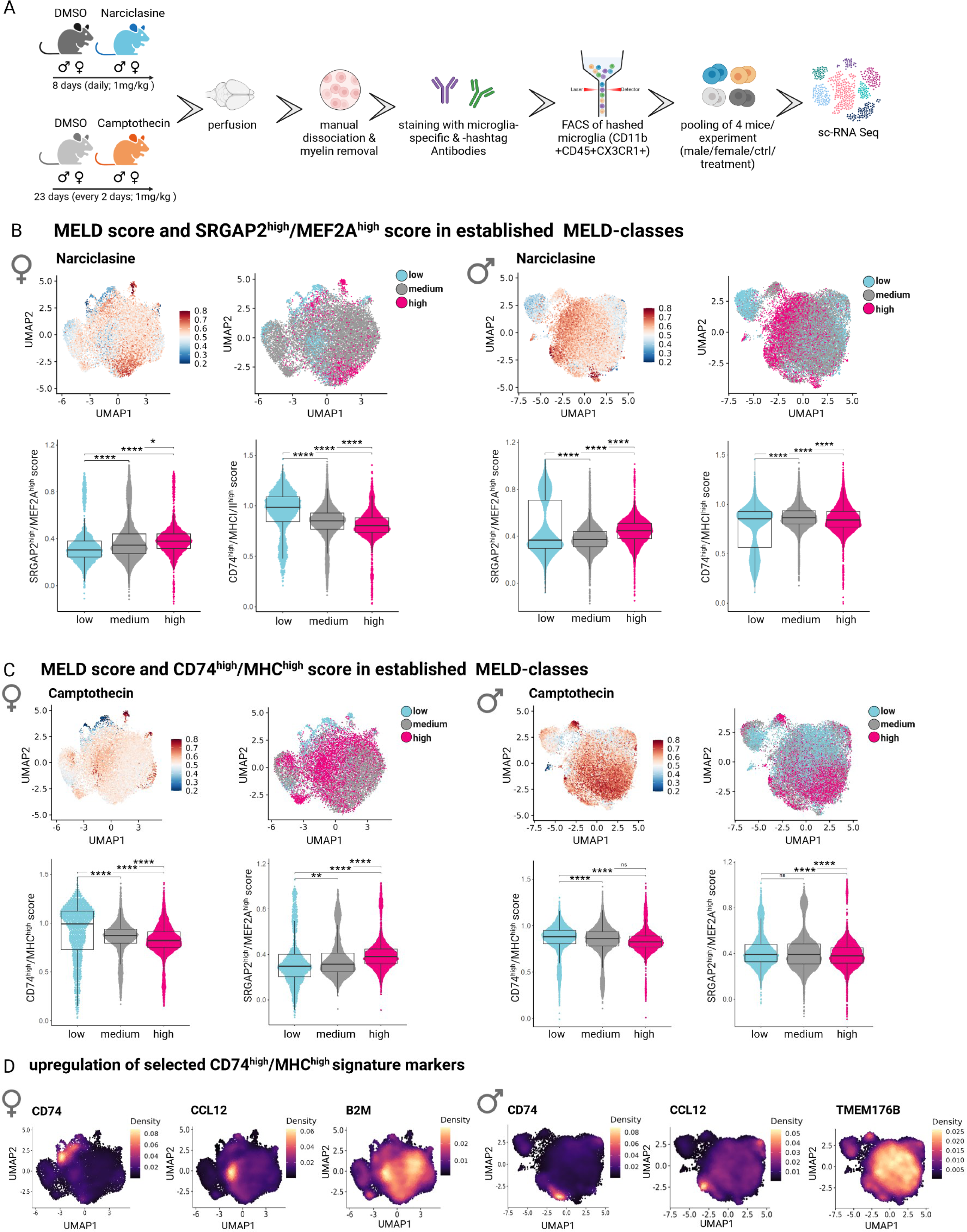
Narciclasine- and Camptothecin-treatment shifts microglial phenotypes *in vivo*. A. Experimental design for Narciclasine- and Camptothecin-treatment of WT mice. For Narciclasine/Camptothecin treatment, 4 female and 4 male WT mice were treated for eight/23 consecutive days with either Narciclasine (1mg/kg)/Camptothecin (1mg/kg) or DMSO as a control via oral gavages. On Day 9/24, perfused brains from one animal/group were sorted by flow cytometry based on CD11b^+^CD45^+^CX3CR1^+^ and subsequently isolated and hashtag- marked microglia from 4 different mice were pooled for sc-RNA Seq using the 10X Chromium platform. **B. MELD score and SRGAP^high^/MEF2A^high^ score in established MELD classes in Narciclasine-treated mice.** For female/male mice, first graph shows the UMAP of all microglia from either female or male mice for all treatment groups (DMSO control for Narciclasine, DMSO control for Camptothecin, Narciclasine treated, Camptothecin treated) with treatment-associated likelihood colored for high likelihood (red) to low likelihood (blue). The second graph shows classification of cells based on treatment-associated likelihood into low (turquoise), medium (grey), high (pink) classes. The lower graphs depict SRGAP^high^/MEF2A^high^ and CD74^high^/MHC^high^ score in each of previously defined treatment-associated likelihood classes. *p.adj ≤ 0.05; **p.adj ≤ 0.01; ***p.adj ≤ 0.001; ****p.adj ≤ 0.0001. **C. MELD score and SRGAP^high^/MEF2A^high^ score in established MELD classes in Camptothecin-treated mice.** For female/male mice, the first graph shows the UMAP of all microglia from either female or male mice for all treatment groups (DMSO control for Narciclasine, DMSO control for Camptothecin, Narciclasine treated, Camptothecin treated) with treatment-associated likelihood colored for high likelihood (red) to low likelihood (blue). The second graph shows classification of cells based on treatment-associated likelihood into low (turquoise), medium (grey), high (pink) classes. The lower graphs depict SRGAP^high^/MEF2A^high^ and CD74^high^/MHC^high^ score in each of the previously defined treatment-associated likelihood classes. *p.adj ≤ 0.05; **p.adj ≤ 0.01; ***p.adj ≤ 0.001; ****p.adj ≤ 0.0001. **D. Upregulation of Top20 CD74^high^/MHC^high^ signature genes in microglia isolated from Camptothecin-treated mice.** Expression level of each of the of Top20 CD74^high^/MHC^high^ signature genes was computed separately and plotted in the respective UMAPs derived from female- or male- compound treated microglia (from lower expression: purple to high expression: yellow). Selected markers are depicted for female microglia on the left side (*CD74*, *CCL12*, *B2M*) and for male microglia on the right side (*CD74*, *CCL12*, *TMEM176B*).

Strikingly, microglia isolated from male and female Narciclasine-treated mice showed a significant upregulation of the SRGAP2^high^/MEF2A^high^ signature across the different classes of treatment-associated likelihood (**Figure 6C**). At the same time, microglia with medium and high Narciclasine treatment-associated likelihood showed a decrease in the CD74^high^/MHC^high^ signature, confirming our data derived from Narciclasine-treated HMC3 cells. These data provide evidence of the capacity of Narciclasine to polarize microglia towards the SRGAP2^high^/MEF2A^high^ signature *in vivo*.

We had some difficulty implementing the published protocol for Camptothecin treatment (see **Methods**), and it is likely that mice were exposed to a low concentration of the compound. The results should therefore be interpreted cautiously. While Camptothecin treatment did not induce the complete CD74^high^/MHC^high^ signature when assessed as a summary score, assessment of the Top 20 marker genes of the CD74^high^/MHC^high^ signature showed upregulation of certain signature genes such as *Ccl12* (both sexes), *Cd74* and *Tmem176b* (males only) and *B2m* (females only) (**Figure 6D**). Further work will be needed to explore the effect of Camptothecin.

### Camptothecin and Narciclasine reduce activated microglia and restore synaptic *in vivo*

We evaluated the effect of our compunds on synaptic density, microglial numbers and morphology in an established amyloidosis model in zebrafish ^39^ (**Figure 7A-B**): purified human Aβ42 peptides are injected via cerebroventricular microinjection (CVMI) into the adult zebrafish brain, mimicking human amyloid deposition, cellular transcriptomic changes and neurodegenerative phenotypes.^39–41^. Consistent with responses to amyloid proteinopathy in humans^42^, this model causes an activation of microglia/macrophages in the zebrafish brain, which can be quantified by morphological analysis of their activation state ^39^. As described for human microglia ^43,44^, amoeboid microglia reflect an activated state (Stage 1 microglia, **Figure 7C**), less round and slightly branched microglia represent a transitionary state (Stage 2 microglia, **Figure 7C**) and slender, branched microglia are defined as resting microglia (Stage 3, **Figure 7C**). Aβ42 also leads to degeneration of synapses and cognitive deficits in the mammalian brain ^45^ and in this amyloid zebrafish model ^39^ where it is marked by loss of the synaptic vesicle protein 2 (SV2) (**Figure 7D**).

**Figure 7.**
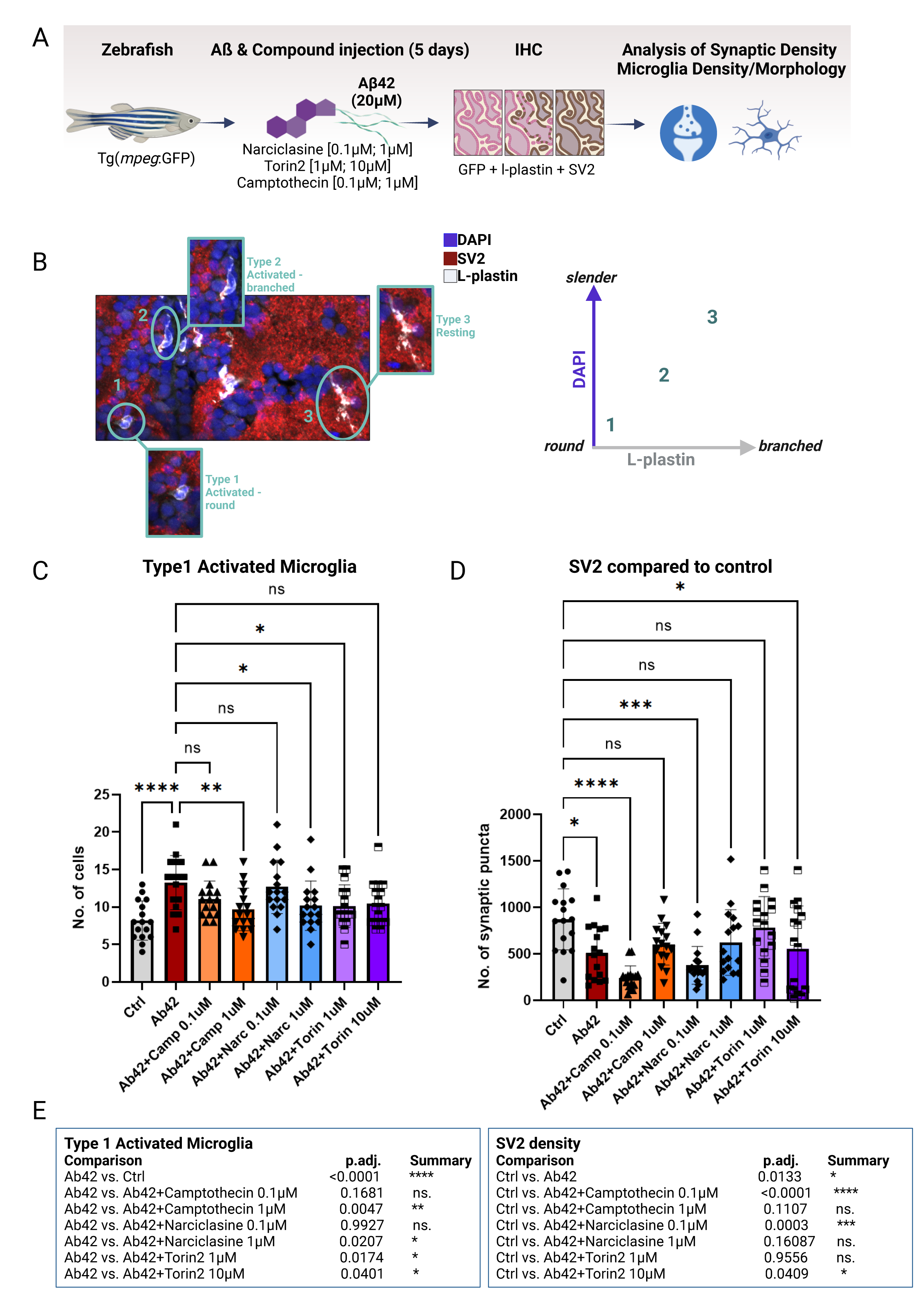
Treatment with Camptothecin, Narciclasine and Torin2 reduces Type 1 activated microglia and restores synaptic density back to control levels in an adult amyloidosis zebrafish model. A. Experimental design of compound-testing in the amyloidosis zebrafish model. Transgenic zebrafish (mpeg:GFP) were injected with 20µM of Aβ42 and co-injected with one dosage of either of the compounds in the brain. After 5 days, zebrafish brains were stained for GFP and L-plastin (microglia), SV2 (synapses) and DAPI (nuclei) to assess microglial density, morphology and synaptic density via confocal microscopy. **B. Overview of morphological classification scheme of microglial activation states.** Microglia were classified into three distinct activation types: Type 1 activated microglia (round-shaped without branching), Type 2 intermediate microglia (activated-branched) , Type 3 resting microglia (slender cell bodies with branching). **C. Quantification of Type 1 activated microglia** (L-plastin; round shaped cell body and missing branches) was performed using confocal images of zebrafish brains harvested 5 days after Aβ42 injection plus compound- or DMSO-injection as control. Bar graphs represent mean cell number ± SD from a total of 16 images/ condition derived from 4 fish/condition. For statistical analysis, 2-way ANOVA followed by Dunnett’s multiple comparison’s test was performed. *p.adj ≤ 0.05; **p.adj ≤ 0.01; ***p.adj ≤ 0.001; ****p.adj ≤ 0.0001. **D. Quantification of synaptic density** (SV2) was performed using confocal images of zebrafish brains harvested 5 days after Aβ42 injection plus compound- or DMSO-injection as control. Bar graphs represent mean cell number ± SD from a total of 16 images/ condition derived from 4 fish/condition. For statistical analysis, 2-way ANOVA followed by Dunnett’s multiple comparison’s test was performed. *p.adj ≤ 0.05; **p.adj ≤ 0.01; ***p.adj ≤ 0.001; ****p.adj ≤ 0.0001. **E. Table summarizing statistical results of quantifications of Type 1 activated microglia and synaptic density.** For statistical analysis, 2-way ANOVA followed by Dunnett’s multiple comparison’s test was performed. *p.adj ≤ 0.05; **p.adj ≤ 0.01; ***p.adj ≤ 0.001; ****p.adj ≤ 0.0001.

Following injection of Aβ42 (20µM) into the zebrafish brain, we injected one low and one higher dose of Narciclasine (0.1µM; 1µM), Torin2 (1µM; 10µM), Camptothecin (0.1µM; 1µM) or DMSO as control (4 fish/group) (see **Methods**). On day 5 post injection, total microglial numbers, microglial morphological activation as well as synapse density (SV2) was quantified (**Figure 7A-B**).

As previously shown in the amyloidosis zebrafish model, the number of activated Stage 1 microglia increased significantly upon Aβ42 injection into the zebrafish brain (^39^; **Figure 7D)**. Camptothecin showed a dose dependent effect in reducing the number of Stage 1 activated microglia, thereby showcasing its potential to modulate microglial activation states in an AD context *in vivo*. Narciclasine and Torin2 also showed a significant decrease in the number of Stage 1 activated microglia (**Figure 7D**). The effect was not dose dependent for Torin2, suggesting that the effect may be saturating. With regards to other microglial activation states, no changes in Type 1 or 3, **Figure S8A**), nor activated-branched microglia could be detected (Type 2, **Figure S7A**).

In sum, all three compounds reduced the amount of Stage 1 microglia, with Camptothecin being the most potent in decreasing Type 1 activated microglia close to the levels of control-treated (DMSO) fish. Whether the observed effects of the drugs can be attributed to direct effects on microglia or whether they are mediated indirectly through the interplay of other CNS-resident cell types remains to be determined.

When assessing synaptic density as previously established^46^, we observed a decrease in synaptic punctae upon treatment with Aβ42 (**Figure 7E**), confirming previously published data from this model system ^39^. Treatment with 1µM Camptothecin led to a restoration of synaptic numbers to levels seen in DMSO-control treated fish. Surprisingly though, the low doses of Camptothecin (0.1µM) and Narciclasine (0.1µM) seemed to reduce the number of synapses even further than Aβ42, which might be explained by the drugs influencing different mechanisms depending on their concentration. Dose-dependent differential mechanisms have been described previously for Camptothecin, with lower doses affecting the inflammatory gene expression while higher doses affect cell proliferation^17,47^. Narciclasine showed a similar pattern of concentration-dependent effect, while Torin2 also restored synaptic punctae to baseline levels. Therefore, both drugs inducing the SRGAP2^high^/MEF2A^high^ signature reduced the amount of Stage 1 activated microglia significantly and restored synapse numbers back to control levels, similar to Camptothecin.

Overall, it is notable that Camptothecin showed the strongest effect in reducing the numbers of Stage 1 activated microglia and also restored the number of synaptic numbers back to control levels (**Figure 7E**). These results are especially interesting in the context of Camptothecin being identified as a drug that is able to mimic the CD74^high^/MHC^high^ cluster and with its selective enhancement of Aβ42-phagocytosis **Figure 4A**). Therefore, our data prioritize Camptothecin and, even more so its FDA-approved analog Topotecan, as the most interesting candidate for modulation of microglial phenotypes in the context of AD pathology.

## Discussion

We have completed a detailed evaluation of a set of tool compounds that emerged from an experimental strategy to develop a chemical toolkit for targeted microglial modulation. Leveraging reference data identifying transcriptionally distinct microglial subtypes^12^, we have developed a pipeline to prioritize, validate and then functionally evaluate candidate compounds that are predicted to engage selected microglial signatures. The three current compounds engage the predicted RNA signatures of two different microglial subtypes, enabling us to link these signatures to empirically measured cellular function, providing a critical link between single cell profiles and behavior of microglial subtypes. The data reported here therefore present a structured functional evaluation of two important transcriptionally defined microglial subtypes - SRGAP2^high^/MEF2A^high^ and CD74^high^/MHC^high^ microglia - in multiple *in vitro* and *in vivo* model systems (summarized in **Table 4**).

One of the most interesting functional results may be the specialization of the CD74^high^/MHC^high^ subtype for enhanced phagocytosis of Aβ, especially in view of the fact that this subtype also suppresses the signature of the SRGAP2^high^/MEF2A^high^ subtype, enriched with susceptibility genes for AD, multiple sclerosis and Parkinson’s disease.

Small molecules and their effect on the modulation of cell fate or cell states have been previously used in other fields such as the stem cell field with the goal of regulating self-renewal, differentiation, trans-differentiation, cell reprogramming or activation for desired therapeutic applications ^48^, but also for direct reprogramming from one cell type to another for transplantation therapy (chemical-compound induced cells – ciCells^49^). Our approach of modulating subtypes of terminally differentiated CNS-resident cells in a rationally designed manner highlights that this approach may be more generally applicable to the brain. The identification of pharmacological compounds that recapitulate established cell-subtype specific transcriptional states allows, on one hand, validation of *in silico* predictions and on the other characterization of the functional properties of human microglial subtypes. One topic of intense debate in the field is the general definition of a cell subtype versus a transient cellular state of reaction^6^. One argument in favor of us successfully modelling microglial subtypes *in vitro* is that for cell subtypes, unique intrinsic features and selective physiological functions should be independent from their microenvironment and thus able to be modulated in model systems such as our 2D and 3D *in vitro* culture systems.

Recently, another approach to model human microglial subtypes *in vitro* has been reported^15^. However, instead of systematically selecting compounds/modulators following a rational design strategy^12^, this effort reversed the approach by exposing iMG to selected CNS- substrates, perturbing their transcriptome. The authors subsequently correlated the induced subtypes with previously described human microglial subtypes derived from snRNAseq datasets ^50^. While this approach is interesting and yielded intriguing insights, it is limited by the variability of the preparation of some of the stimulants (such as synaptic preparations) and the lack of direct translation to therapeutic development. Our structured pharmacological approach arguably bears higher potential to yield reproducible results when applied to model systems, results: in our case, translation to at least two *in vivo* systems and repurposing of Food and Drug Administration (FDA) approved compounds such as Topotecan.

Another aspect of our manuscript that bears highlighting is the mitochondrial phenotyping of human microglial subtypes, establishing a connection between specific mitochondrial phenotypes (or mitotypes) in relation to a defined microglial subset. Immuno-metabolic differences have been reported among circulating human leukocytes^36^. Here we relate specific microglial subsets to mitochondrial phenotypes in primary human microglia and model systems. We confirm an initially defined mitotype of reduced oxidative phosphorylation for MEF2A^high^/SRAGP2^high^ microglia in the human microglial cell line HMC3 as well as in microglia isolated and identified from female and male Narciclasine-treated mice (**Figure S7B**).

Further, our mitotyping analysis revealed an enrichment of CD74^high^/MHC^high^ microglia in iron homeostasis, metals and cofactors as well as ROS and glutathione metabolism in freshly isolated human microglia and Camptothecin-treated HMC3 microglia (**Figure 6**). These findings are highly interesting with regards to the described CD74^high^/MHC^high^ cluster showing enhanced Aβ uptake (**Figure 5B**). While a clear relation between accumulation of free iron in the brain parenchyma and neurodegenerative diseases has been reported ^51^, microglia have been shown to store iron-bound ferritin, and iron-overload causes marked shifts in the transcriptional states of microglia^52^, potentially indicating a shift in microglial subtype identity.

As a product of our study, we provide to the microglial community an initial set of compounds with which to stock a pharmacological toolbox for targeted immunomodulation. This initial characterization of the functional consequences of our compounds also serves as a proof of principle for our approach to model specific human microglial subsets *in vitro* to study their function. Multiple studies by colleagues ^53,54^ and our laboratory^11,12^ have repeatedly identified a neuroimmunologically active human microglial subpopulation characterized by CD74^high^ expression (here: CD74^high^/MHC^high^) which we can now mimic using low-dosages of the Topoisomerase inhibitors Camptothecin or Topotecan. We started characterizing several functions of our CD74^high^/MHC^high^ model system and found striking results with regards to its function in Aβ phagocytosis (**Figure 5B**), anti-inflammatory cytokine secretion (**Figure 5F**) and its potential to restore activated microglia to more morphologically quiescent state and protect synaptic density in a zebrafish amyloid proteinopathy model (**Figure 8D-E**). Our current functional analyses are initially conducted using the human embryonic microglia (HMC3) cell line which has the usual limitations of an immortalized cell line but recapitulates many features of myeloid and microglial cell behavior. We address this limitation by validating our work in iMG and *in vivo* model systems; these results lay the groundwork for further functional characterization of our compounds in a variety of complementary model systems.

We identified Topotecan as a structural and functional analog of Camptothecin that robustly induces the CD74^high^/MHC^high^ signature and functionally mimics Camptothecin’s downregulation of pro-inflammatory cytokine secretion in HMC3 cells. These results are of particular interest as Topotecan is an FDA-approved drug, originally authorized for the treatment of small-cell lung cancer and cervical cancer ^16^. Topotecan is clinically more effective and safer than Camptothecin ^55^ and in the light of our findings, Topotecan might represent an entry point for therapeutic development of a microglia-targeted AD-therapy with the potential to shift microglial subpopulations towards the reduced CD74^high^/MHC^high^ phenotype that enhances Aβ uptake. Thus, this class of compounds could have a role early in the trajectory to AD, during the initial phase of accumulation of amyloid proteinopathy; it may help to boost the effects of current approved anti-amyloid antibodies ^56,57^ since the mechanisms are different.

Alternatively, it can be an alternative for those who do not respond to or have adverse events from antibody treatments. Side effects of current clinical doses in cancer therapy using Topotecan, e.g. hematopoietic toxicity, are quite serious due to its function as a Topoisomerase I (TopI) inhibitor. Its anti-inflammatory effects on microglia, however, are ascribed to lower doses in preclinical studies of Parkinson’s disease (PD^30^) and our own dataset. Thus, as with repurposing of chemotherapy agents such as cyclophosphamide and cladribine in multiple sclerosis ^58,59^, initial clinical research in AD could start off with low Topotecan doses followed by TSPO PET imaging and other markers of microglial activation in CSF.

Additionally, discussions regarding a new generation of Camptothecin analogs with minimal TopI inhibitory activity and maximal Camptothecin-based structural efficacy are currently under way ^55^. This new generation of Camptothecin-analogs might also bear great potential for our proposed microglia-targeting approaches in AD therapy and potentially those of other neurodegenerative diseases, as the goal of therapy probably involves subtle shifts in the frequency of certain microglial subtypes and not wholesale elimination of microglia.

Overall, this deep characterization of three putative tool compounds for microglial modulation has validated our approach to identify chemical matter that influence microglia in a specific manner. Its foundation on a large dataset of single cell transcriptomes from live human microglia ^12^ is probably an important component of its success. The three compounds expand the list of reagents with which to manipulate human microglia, and we look forward to colleagues elaborating this initial characterization in other contexts as the community continues to build its toolkit of translational reagents. At this time, the case for further investigation of topoisomerase inhibition and other effects of Camptothecin and Topotecan in AD and other neurologic diseases is compelling and will hopefully contribute to preventing these diseases.

## Acknowledgements

The work was supported by the Chan-Zuckerberg Initiative’s Neurodegeneration Challenge Network grant CS-02018-191971. Some of the work also emerged from support from NIH/NIA grants R01 AG070438, U01 AG061356, RF1 AG057473, R01AG048015. Research reported in this publication was supported by the National Institute of General Medical Sciences of the National Institutes of Health under Award Number T32GM007367 and by the National Cancer Institute of the National Institutes of Health under Award Number F30CA261090.

Research reported in this publication was partially performed in the Columbia Center for Translational Immunology and P&S Flow Cytometry Core, the Columbia Stem Cell Initiative Flow Cytometry Core at Columbia University and the Proteomics and Macromolecular Crystallography Core in the Columbia University Herbert Irving Comprehensive Cancer Center. AAS is supported by The Thompson Foundation (TAME-AD) and the Henry and Marilyn Taub Foundation. We further thank the New York Genome Center for a long-standing collaboration and efforts in RNA-Sequencing for the study. In addition we thank the Zebrafish International Resource Center (ZIRC) for providing the Tg (mpeg1:EGFP) zebrafish line.

All illustrations were created with BioRender.com.

## Supplemental tables

Table S1: Statistical analysis supporting Supplementary Figure S1C. Expression levels of different expression modules of human microglial subpopulations across compound-treated HMC3 microglial subclusters.

Table S2: Human MitoCarta3.0 list and Maestro rankings of mitochondrial localization for all human genes ^37^.

## Methods

### In vitro sc-RNA Seq datasets

#### Single-cell RNA Sequencing of compound-treated HMC3s

0.5x10^6^ HMC3 microglial cells were seeded into a 6-well plate and incubated o.n. The next day, microglia were treated with the respective concentrations of Camptothecin (1µM; EMD Millipore; Cat #:390238), Narciclasine (0.1µM; Millipore Sigma; Cat #: SML2805), Torin2 (10µM; Cayman Chemical Company; Cat #: 14185) or DMSO (Sigma-Aldrich, Cat #:472301) as control and incubated for 24hrs before harvest. Cells were trypsinized (Gen Clone; Cat #:25-510F) and each treated HMC3 sample was subsequently barcoded using “hashing antibodies” consisting of antibodies targeting beta-2-microglobulin and CD298 and conjugated to “hash-tag” oligonucleotides (HTOs). In order to do so, cells were washed with PBS (Corning; Cat#:21-040-CV), centrifuged at 300g, 4°C, 5min and filtered through a 70µm filter (fisher scientific; Cat#: 08-771-23). Following centrifugation, cells were resuspended in 25µl PBS+0.04% BSA (pluriSelect; Cat#: 60-00020-10BSA), 1µl of FcBlock (BioLegend, Cat#: 422301) was added and cells were incubated for 10min on ice. 2ul of each hashtag antibody (TotalSeq™-B0256 anti-human Hashtag 6 Antibody, Cat#: 394641; TotalSeq™-B0257 anti-human Hashtag 7 Antibody, Cat#: 394643; TotalSeq™-B0258 anti-human Hashtag 8 Antibody, Cat#: 394645; TotalSeq™-B0259 anti-human Hashtag 9 Antibody, Cat#: 394647 ;0.5 mg/mL) was mixed with 25µl PBS+0.04% BSA, passed through 0.1um filter at 12000g for 2 min, 4°C and antibody mix for each hashtag was added to the respective tube containing cells in FcBlock. Cells were incubated for 30mins on ice and subsequently washed 4 times with staining buffer. Cells for each sample were then counted, mixed 1:1:1:1, filtered, then counted using c) and subsequently subjected to single-cell library preparation.

The single-cell library preparation was constructed using 10x Chromium Next GEM Single Cell 5’ Reagent Kits v2 (Dual Index) with Feature Barcode technology for Cell Surface Protein (10x Genomics, Pleasanton, CA) according to the manufacturer’s protocol. Briefly, a total of 30,000–40,000 cells were loaded on the 10x Genomics chromium controller single-cell instrument. Reverse transcription reagents, barcoded gel beads, and partitioning oil were mixed with the cells for generating single-cell gel beads in emulsions (GEM). After reverse transcription reaction, the GEMs were broken. Feature Barcode cDNA amplification was performed. The amplified cDNA was then separated by SPRI size selection into cDNA fractions containing mRNA derived cDNA (>400bp) and HTO-derived cDNAs (<180bp), which were further purified by additional rounds of SPRI selection. Independent sequencing libraries were generated from the mRNA and HTO cDNA fractions, which were analyzed and quantified using TapeStation D5000 screening tapes (Agilent, Santa Clara, CA) and Qubit HS DNA quantification kit (Thermo Fisher Scientific). Libraries were pooled and sequenced together on a NovaSeq 6000 with S4 flow cell (Illumina, San Diego, CA) using paired-end, dual-index sequencing with 28 cycles for read 1, 10 cycles for i7 index, 10 cycles for i5 index, and 90 cycles for read 2.

#### Analysis of single-cell RNA Sequencing of compound-treated HMC3s

FASTQ files of single-cell sequencing libraries were processed using the “count” command of Cell Ranger (10x Genomics, version 6.0.1). Gene expression library and hashtag oligo library were processed together, and the human transcriptome (GRCh38-2020-A) was used as the reference for alignment. As HMC3s have higher median feature number as compared to live human microglia, our filtering thresholds were adjusted and more aggressive doublet removal approaches were applied. We did not set a top bound for UMI-based filtering, only discarding low-quality cells below 500 UMIs. Hashtags were deconvoluted with demuxmix. Subsequently, we checked for doublets by using a recently published tool, DoubletFinder ^60^, that was the top performer in a recent benchmark of computational doublet detection approaches ^61^. We followed the standard workflow for DoubletFinder analysis, setting our expected rate to 0.2 based on extrapolation of expected multiplet rates based on loading densities from the 10x Genomics website. The parameters we used for DoubletFinder were: 15 PCs and pN of 0.25. pK was calculated from our dataset. Substantial overlap was found between the doublets identified by demuxlet and DoubletFinder, and DoubletFinder-identified doublets generally had higher counts than identified singlets. As such, all doublets from demuxmix and DoubletFinder were excluded from further analysis. Clustering was conducted with the standard single-sample SCTransform workflow. Briefly, SCTransform was run with default settings, followed by PCA calculation with RunPCA. We chose to retain 20 PCs for downstream analysis by inspecting an elbow plot. To optimize clustering resolution, the recently reported method ChooseR ^62^, which compares silhouette scores across separate resolutions of Louvain clustering, was used. The final choice of resolution was 0.6, and this was used for downstream analyses. Differentially expressed genes per cluster were calculated as detailed in ^12^ . Annotation of the genes associated with different clusters was performed by way of Reactome pathway annotation using the clusterprofiler package, as described in ^63^. To calculate module enrichment per cell, the Seurat AddModuleScore function was used. In brief, this function takes an input list of genes, and for each of these genes, assigns a “background” of 100 genes expressed at similar levels, then calculates the overall enrichment of input genes versus control gene sets on a per-cell level ^64^. This approach was applied for the top 100 genes per cluster to derive a cluster enrichment score on a per-cell level. Testing for significant differences between clusters in module scores was performed with pairwise Wilcoxon testing. Assessment of enrichment of cluster signatures in association with compound treatment, in contrast, was performed with GSEA ^65^. Here, log fold change was calculated for all genes in the dataset between control (both, DMSO and untreated) and the drug treatment of interest, and genes were ranked based on average log fold change across all cells in each condition. Enrichment of cluster signatures in this rank-ordered list was then performed with GSEA.

#### Evaluating mitotype enrichment in transcriptomic data

Using mitochondrial pathways defined from the MitoCarta3.0 gene sets for murine and human datasets ^37^, we evaluated enrichment of different sets in the rank-ordered log-fold-change lists for compound-treated versus control/DMSO conditions for either bulk RNA-seq or for aggregated cells in scRNA-seq. GSEA was used as previously described to calculate enrichment of genes associated with different mitochondrial pathways in sets of genes associated with different compound treatment conditions.

#### Mitochondrial pathway analysis in human single-cell sequencing data

To delve more into the metabolic differences between compound treatment conditions in the human single-cell data, we leveraged recent annotated gene sets from MitoCarta3.0 ^37^ as mentioned in **Evaluating mitotype enrichment in transcriptomic data**. Using gene sets tied to distinct mitochondrial properties, we calculated enrichment of different sets in upregulated cluster signatures using a hypergeometric test with an FDR-corrected threshold for significance at *q* = 0.01 ^66^. The results of this analysis were visualized in heatmaps where the color intensity corresponds to the -log10 p-value of the FDR q-value for enrichment of each given gene set in the differentially expressed genes associated with a given cluster.

FA000010 (RUCDR/BiologyX) iPSCs were differentiated into hematopoietic precursors cells (HPCs) using the STEMdiff Hematopoietic kit (STEMCELL Technologies) largely by manufacturer’s instructions. In brief, on day -1 iPSCs were detached with ReLeSR and passaged to achieve a density of 1–2 aggregates/cm^2^ of 100-150 cells. Multiple densities were plated in parallel. On day 0, colonies of appropriate density were switched to Medium A from the STEMdiff Hematopoietic Kit to initiate HPC differentiation. On day 3, cells were switched to Medium B with a full media change and fed again with a full media changed on day 5. Cells remained in Medium B for the rest of the HPC differentiation period with Medium B overlay feeds every other day. HPCs were collected 3 independent times by gently removing the floating population with a serological pipette at days 11, 13 and 15 (or days 12, 14 and 16). HPCs were either cryobanked in 45% Medium B, 45% knockout serum replacement (ThermoFisher) and 10% DMSO and stored in liquid nitrogen or directly plated for iMGL induction. HPCs were terminally differentiated at 28,000-35,000 cells/cm^2^ in microglia medium (DMEM/F12, 2X insulin-transferrin-selenite, 2X B27, 0.5X N2, 1X glutamax, 1X non-essential amino acids, 400 mM monothioglycerol, and 5 mg/mL human insulin (ThermoFisher)) freshly supplemented with 100 ng/mL IL-34, 25 ng/mL M-CSF (R&D System) and 50 ng/mL TGFβ1 (STEMCELL Technologies) for every other day until day 24. On day 25, 100 ng/mL CD200 (Bon Opus Biosciences) and 100 ng/mL CX3CL1 (R&D Systems) were added to Microglia medium to mimic a brain-like environment. For all experiments in this study, iMGLs were treated between day 28 to 29. On day 28/29 of iMG differentiation, iMGs in 12-wells were treated for 24 hours with the following compounds and the following concentrations:

Differentiated microglia were cultured in 500µl microglia medium (DMEM/F12, 2X insulin-transferrin-selenite, 2X B27, 0.5X N2, 1X glutamax, 1X non-essential amino acids, 400 mM monothioglycerol, and 5 mg/mL human insulin (ThermoFisher)). For compound treatment, 2x solutions were prepared for each compound and subsequently 500µl of 2x solutions was added to each well. After 24hrs, treated iMGs were harvested in low-protein binding tubes (fisher scientific; Cat#: 13-864-407) using media and PBS (Corning; Cat#:21-040-CV) to wash off iMGs from cell culture wells. Subsequently, cells were washed 1x with PBS, centrifuged at 300g, 4°C, 5min, filtered through a blue lid filter (FACS tube), resuspended in 100uL PBS+0.04% BSA for counting using counting chambers (Bulldog Bio, Portsmouth, NH) and subsequently subjected to single-cell library preparation.

The single-cell library preparation was constructed using 10X Chromium Next GEM Single Cell 3’ Reagent Kits v3.1 (Dual Index) according to the manufacturer’s protocol. Briefly, a total of ∼10,000 cells were loaded on the 10X genomics chromium controller single-cell instrument. Reverse transcription reagents, barcoded gel beads, and partitioning oil were mixed with the cells for generating single-cell gel beads in emulsions (GEM). After reverse transcription reaction, the GEMs were broken. cDNA amplification was performed. The amplified cDNA was then separated by SPRI size selection into cDNA fractions containing mRNA derived cDNA (>300bp) which were further purified by additional rounds of SPRI selection. Sequencing libraries were generated from the mRNA cDNA fractions, which were analyzed and quantified using TapeStation D5000 screening tapes (Agilent, Santa Clara, CA) and Qubit HS DNA quantification kit (Thermo Fisher Scientific).

#### Pre-processing of single-cell RNA Sequencing of compound-treated iPSC-derived microglia

The h5 file output by Cell Ranger for each sample was read in R (4.2.2) using the Seurat package (4.3.0). For quality control, we removed cells with a mitochondrial RNA percentage above a statistical threshold of either two absolute deviations above the median mitochondrial reads within the sample, or 1.5 times the interquartile range above the third quartile, whichever is a lower threshold. If this threshold is lower than 10%, we set the threshold to 10%. All ribosomal genes, mitochondrial genes and pseudogenes were removed. Cells with more than 500 UMI count were kept for downstream analysis. To predict and remove potential doublets, we applied DoubletFinder (2.0.3) ^60^ on the first 15 principal components of the iMG data, using the function *doubletFinder_v3* (with pN=0.25, nExp = 0.075*(# cells in library)*(1- homotypic proportion), sct = F). The homotypic proportion was estimated using the *modelHomotypic* function, whereas the pK parameter was optimized for each sample using *paramSweep_v3.* To correct for batch effect in the iMG data, we used the scVI model from the scvi-tools package in python (3.8). scVI generates non-negative corrected counts and has recently been benchmarked to perform well in data integration ^69^. We trained the scVI model using raw counts as input, with two hidden layers (n_layers = 2), 30 dimensions of the latent space (n_latent = 30) and assuming a negative binomial gene count distribution. The batch corrected counts were then log normalized using the *log1p* function from the numpy package (1.21.6). The scVI generated latent space were then used for dimension reduction using Uniform Manifold Approximation and Projection (UMAP). The batch corrected counts results from scVI were used for query mapping and MELD analysis described below.

#### Single-cell RNA Sequencing of compound-treated cerebral organoids with incorporated iPSC-derived microglia

Cortical brain organoids (CO) were cultured using the STEMdiff^TM^ Cerebral Organoid Kit (#08570, #08571). This four-stage protocol is based on the original Lancaster protocol ^70^. hiPSC lines used were MSN38 (Mount Sinai Normal, Icahn School of Medicine at Mount Sinai) and WTC-11 (Allen Institute for Cell Science). At mature age the COs were co-cultured with microglial progenitors (derived from WTC-11) in order to obtain microglia-containing organoids (CO-iMG), as previously published ^21^. For more details on the co-culture protocol, please refer to Buonfiglioli et al. *in preparation*. On day 6/7 of co-culturing iPSC-derived microglia (iMGs) with the respective cerebral organoids derived from the same lines (CO-iMGs) were treated in 500µl CO-iMG-specific media containing compounds of interest (Camptothecin (EMD Millipore; Cat #:390238), Narciclasine (Millipore Sigma; Cat #: SML2805), Torin2 (Cayman Chemical Company; Cat #: 14185) or DMSO as control (Sigma-Aldrich, Cat #:472301). 2x compound-containing media was prepared and for treatment, 250µl were removed from the 500µl-containing 24-well plate and subsequently 250µl of the 2x solution were added to achieve 1x solution for compound treatment. Final treatment concentrations for the compounds were used as follows: Camptothecin 0.1µM and 0.5µM, Narciclasine 0.01µM, Torin2 10nM. CO-iMGs were incubated for 24hrs at 37°C, 5%CO2. Subsequently, cerebral organoids were collected, transferred into a tissue homogenizer (5ml) containing 2ml CO-iMG media, and dissociated on ice using the loose pestle. Cell solution was subsequently filtered through a 35µm strainer (5ml, fisher scientific, Cat#: 08-771-23) into a low protein binding tube (fisher scientific; Cat#: 13-864-407). Cell solution was centrifuged at 300g, 4°C, 5min, twice washed with 1x with PBS (Corning; Cat#:21-040-CV), filtered, centrifuged at 300g, 4°C, 5min and resuspended in 60uL of PBS (Corning; Cat#:21-040-CV) containing 0.04% BSA (pluriSelect; Cat#: 60-00020-10) for counting using disposable counting chambers (Bulldog Bio, Portsmouth, NH). Subsequently, volumes corresponding to the same cell number of corresponding samples (DMSO, Camptothecin, Narciclasine, Torin2) from each genetic line (MSN38 or WTC11) were mixed 1:1 to achieve equal cell loading for each sample within one single-cell sequencing experiment.

For the 10x Genomics, we targeted 30,000 cells for cell loading. The single-cell library preparation was constructed using 10X Chromium Next GEM Single Cell 3’ Reagent Kits v3.1 (Dual Index) according to the manufacturer’s protocol. Briefly, a total of ∼30,000 cells were loaded on the 10X genomics chromium controller single-cell instrument. Reverse transcription reagents, barcoded gel beads, and partitioning oil were mixed with the cells for generating single-cell gel beads in emulsions (GEM). After reverse transcription reaction, the GEMs were broken. cDNA amplification was performed. The amplified cDNA was then separated by SPRI size selection into cDNA fractions containing mRNA derived cDNA (>300bp) which were further purified by additional rounds of SPRI selection. Sequencing libraries were generated from the mRNA cDNA fractions, which were analyzed and quantified using TapeStation D5000 screening tapes (Agilent, Santa Clara, CA) and Qubit HS DNA quantification kit (Thermo Fisher Scientific).

#### Preprocessing of single-cell RNA Sequencing of compound-treated cerebral organoids with incorporated iPSC-derived microglia

To demultiplex the samples, cells were grouped into two clusters based on SNPs in RNA-seq reads using freemuxlet software (https://github.com/statgen/popscle). Positions of SNPs were obtained from a VCF file of the 1000 Genomes Project phase 3 after excluding SNPs with minor allele frequency < 10%. Average expression of *XIST* RNA was computed in each cluster. A cluster with higher *XIST* expression should be female origin and thus was annotated as MSN38. Conversely, a cluster with lower *XIST* expression was annotated as WTC11. Quality control (i.e., low quality cell removal, gene filtering and doublet removal) was performed as in the iPSC-derived microglia preprocessing. To identify microglia from the organoid data, we performed the following standard Seurat pipeline to cluster cells: *SCTransform*, *RunPCA*, *RunUMAP* (using the first 30 principal components with default parameters), *FindNeighbors,* and *FindCluster* using a resolution of 0.5. Cells of the cluster with the highest co-expression of microglial markers (*AIF1, C1QA, C1QB, CSF1R, CTSS, CX3CR1, GPR34, HEXB, ITGAM, MERTK, P2RX7, P2RY12, PTPRC, SLC2A5, TGFBR1, TMEM119*) were annotated as microglia. This process resulted in 492 cells annotated as microglia. Here we focused on DMSO- (n = 131) and Camptothecin-treated cells (n = 130) for further analysis, since these two conditions contained a sufficient number of cells for analysis.

To understand the similarity and differences between *in vitro* iMGs and organoid-iMGs in drug response, we mapped the CO-iMGs treated with DMSO and Camptothecin as query onto the iMGs, of which the batch-corrected count matrix was used as reference data. The reference-query mapping was performed using the Seurat query mapping pipeline ^22^. Briefly, the query data was first normalized using the function *NormalizeData* (with normalization.method = “LogNormalize” and scale.factor = 10000). We then identified the top 1000 variable genes from the log-normalized query data and from the log-transformed scVI batch-corrected counts of the reference data, using the *FindVariableFeatures* (with nfeatures = 1000). A set of mapping anchors (pairs of cells that encode the relationships between the two datasets) between the reference and query was identified and scored using the *FindTransferAnchors* function (using the first 15 principal components with default parameters). The anchors were then used as input for the *MapQuery* function to calculate the transformation vectors that transform query gene expression to be combined with the reference for downstream analyses, including UMAP dimension reduction and MELD analysis.

### Treatment of C57BL/*6J* mice with Narciclasine and Camptothecin

#### Mice

12-18 weeks old male and female C57BL6/N mice (The Jackson Laboratory, United States) were used throughout the study. 3–5 animals were kept in individually ventilated cages (IVCs). The animal vivarium was a specific-pathogen-free (SPF) holding room, which was temperature- and humidity-controlled (20-26°C, 30-70% humidity as per the Guide for Care and Use of Laboratory Animals) and kept under a light–dark cycle (lights off: 07:00 AM–07:00 PM). All animals had ad libitum access to standard rodent chow ( PicoLab 5053 rodent diet 20; LabDiet, Cat#: 3005740-220) and water throughout the entire study. All procedures described in the present study were approved by and complied with the guidelines of the Institutional Animal Care and Use Committee of the Nathan Kline Institute and Columbia University Medical Center (Animal Care Protocol AC-AABR4602 (Y1 M05).

#### Narciclasine treatment

Narciclasine treatment protocol was adapted from a previous publication assessing the effects of oral gavages of Narciclasine on the murine CNS ^38^. Narciclasine (R&D Systems, Cas#: 29477-83-6) was dissolved in 5% DMSO (Tocris Bioscience, Cat#: 3176; Batch: 68A) containing PBS (Corning; Cat#:21-040-CV) to a stock concentration of 0.1mg/ml. Subsequently, Narciclasine stock was stored in small aliquots at -20°C until treatment. As a treatment control, a 5% DMSO (Tocris Bioscience, Cat#: 3176; Batch: 68A) containing PBS (Corning; Cat#:21-040-CV) solution was applied by oral gavages. Mice were treated with Narciclasine at a concentration of 1mg/kg through oral gavages for eight consecutive days and sacrificed on Day 9. Each treatment group (Control females, Control males, Narciclasine females, Narciclasine males) consisted of five mice. Treatments were conducted in 5 cohorts in total, each one consisting of one control female, one control male, one Narciclasine female, one Narciclasine male and cohorts were started in intervals of 1-6 days. For Volume of oral gavages was based on the weight acquired on Day1 of treatment and weight was monitored and recorded on a daily basis using a digital scale.

#### Camptothecin treatment

Camptothecin treatment protocol was adapted from a previous publication assessing the effects of oral gavages of Camptothecin on neuroprotection ^30^. Camptothecin (Millipore Sigma, Cat#: 390238- 25MG, Lot#: 3837159) was dissolved in DMSO (Tocris Bioscience, Cat#: 3176; Batch: 68A) to a stock concentration of 1mg/ml. Subsequently, Camptothecin stock was stored in small aliquots at -20°C until treatment. For Camptothecin treatment, a 0.3mg/ml Camptothecin solution was prepared freshly every day of treatment by diluting the 1mg/ml stock solution 1:3 in PBS (Corning; Cat#:21-040-CV). For control mice, at the day of treatment, DMSO (Tocris Bioscience, Cat#: 3176; Batch: 68A) was diluted 1:3 in PBS (Corning; Cat#:21- 040-CV). Mice were treated at a concentration of 1mg/kg Camptothecin every second day for a total of 23 days and sacrificed at Day 24. Each treatment group (Camptothecin females, Camptothecin males, Camptothecin females, Camptothecin males) consisted of five mice. Each treatment group (Control females, Control males, Camptothecin females, Camptothecin males) consisted of five mice. Treatments were conducted in 5 cohorts in total, each one consisting of one control female, one control male, one Camptothecin female, one Camptothecin male and cohorts were started in intervals of 2-4 days. Volume of oral gavages was based on the weight acquired on Day1 of treatment and weight was monitored and recorded on a daily basis using a digital scale.

### Isolation of microglia and single-cell RNA Sequencing from compound-treated C57BL/*6J* mice

#### Preparation of mouse brains for microglia isolation

Narciclasine- or Camptothecin treated mice were sacrificed on Day 9, respectively Day 24, perfused intracardially with 15 mL cold PBS (Corning; Cat#:21-040-CV) via a 20ml syringe (Millipore Sigma, Cat#: Z116882) with a 23G needle (Millipore Sigma, Cat#: CAD9931) and brains were removed and stored in Hibernate A media (Gibco, Cat#: A1247501) on ice. For each individual mouse, cerebellum and olfactory bulb were removed and the right hemisphere was further processed for single-cell sequencing experiments. Brain samples were dissociated and prepared for further processing using the brain cell isolation protocol published by ^71^. In short, each hemisphere was minced in 1ml Hibernate A media (Gibco, Cat#: A1247501) on a petri-dish (Corning, Cat#: CLS430167) using a scalpel (Sigma Millipore, Cat#: S2646) before transferring the sample to a 1.5 ml Hibernate A media (Gibco, Cat#: A1247501) containing 1ml Dounce Homogenizer using the loose pestle (Gibco, Cat#: A1247501). Homogenized brain lysate was then transferred to a 15ml Falcon tube (Sigma Aldrich, Cat#: CLS430052) by filtering through a 70µM strainer (Millipore Sigma, Cat#: CLS431751) and centrifuged for 10min at 4°C and 350g. Subsequently, the pellet was resuspended in 1ml PBS (Corning; Cat#:21- 040-CV), 500µl isotonic Percoll was added (Sigma Aldrich, Cat#: GE17-0891-02) and the volume was adjusted to 2ml using PBS (Corning; Cat#:21-040-CV) if needed. 2ml of PBS were then carefully layered on top of the Percoll solution and layered samples were centrifuged at 3000g for 10mins at 4°C. Myelin ring and upper phase were removed and leftover cellular pellet was washed using PBS (Corning; Cat#: 21-040-CV) followed by centrifugation at 400g for 10mins at 4°C. Subsequently, cells were stained with hashtags and cell surface markers for isolation using flow cytometry and further downstream processing for single-cell sequencing experiments. In detail, cells were resuspended in PBS (Corning; Cat#:21-040-CV), transferred to a low-protein binding tube (fisher scientific; Cat#: 13-864-407), washed again in PBS (Corning; Cat#:21-040-CV) and resuspended in 25µL PBS+0.04% BSA (pluriSelect; Cat#: 60- 00020-10). 0.25µL of TruStainFcX (BioLegend, Cat#: 101319) were added and samples were incubated for 10mins on ice. For each processed cohort consisting of Control females, Control males, and compound-treated females, compound-treated males, each treatment group sample was subsequently stained with a specific hashtag as every cohort processed on the same day was single-cell sequenced in one single-cell sequencing experiment. To 4µl of each hashtag antibody (Hashtag B 301#, 0.5 mg/ml (BioLegend, Cat#: 155831); Hashtag B 302# 0.5 mg/ml (BioLegend, Cat#: 155833); Hashtag B 303# 0.5 mg/ml (BioLegend, Cat#: 155835); Hashtag B 304# 0.5 mg/ml (BioLegend, Cat#: 155837)) 25uL of PBS+0.04% BSA (pluriSelect; Cat#: 60-00020-10) were added and the solution passed through 0.1um filter (fisher scientific, Cat#: 10439753) at 12000g for 2 min, 4°C. At the same time, flow cytometry staining master mix was prepared containing 1:200 dilutions of PE anti-mouse/human CD11b Antibody (BioLegend, Cat#:101208, clone M1/70), APC anti-mouse CX3CR1 (Antibody BioLegend, Cat#: 149008, clone SA011F11), APC/Cyanine7 anti-mouse CD45.2 Antibody (BioLegend, Cat#: 109823, clone 104) in 50µl cell staining buffer/sample (BioLegend, Cat#: 420201). After the 10min incubation with TruStainFcX, antibody master mix was added to the cells and solution incubated for 30mins on ice, followed by 4x washed using PBS+0.04% BSA (pluriSelect; Cat#: 60-00020-10) using centrifugation for 5mins at 300g at 4°C. Finally, cells were resuspended in 1ml cell staining buffer and 1µl SYTOX Blue (ThermoFisher, Cat#: S34857) for live/dead staining was added followed by 5-10mins incubation at RT.

#### Sorting of microglia from mouse brain using flow cytometry-activated cell sorting (FACS)

Microglia were sorted from stained brain lysate using a Sony MA900 Multi-application cell sorter (Sony, Cat#: ) by selecting brain cells using SSC-A vs. FSC-A gating, followed by exclusion of duplet cells using FSC-H vs. FSC-A gating, followed by selection of live cells by selecting SYTOX Blue-negative cells in a FSC.H vs. SYTOX-Blue -A plot. From live cells, microglia were subsequently gated and sorted as CD11b- and CD45-double positive cells (CD45.2-APC/Cy7-A vs. CD11b-PE-A) and further gated for CXC3CR1-positive cells (CX3CR1-APC-A vs. CD11b-PE-A). CD11b+CD45.2+CX3CR1+ microglia were sorted into a pre-cooled 5ml low-protein binding falcon tube (fisher scientific; Cat#: 13-864-407) containing 0.5ml cell staining buffer (BioLegend, Cat#: 420201). Walls of the tubes were washed and cells pelleted at 300g, 5mins, 4°C. For counting, cells were resuspended in 60-100µl, and subsequently, all four samples from each cohort mixed 1:1:1:1, pelleted, resuspended in a final volume of 100µl, filtered through a 35µM strainer (5ml, Fisher Scientific, Cat#: 08-771-23) and subjected to library preparation.

#### Preparation of library for RNA-Seq experiments

The single-cell library preparation was constructed using 10x Chromium Next GEM Single Cell 3’ Reagent Kits v3.1 (Dual Index) with Feature Barcode technology for Cell Surface Protein (10x Genomics, Pleasanton, CA) according to the manufacturer’s protocol. Briefly, a total of 30,000 cells were loaded on the 10x Genomics chromium controller single-cell instrument. Reverse transcription reagents, barcoded gel beads, and partitioning oil were mixed with the cells for generating single-cell gel beads in emulsions (GEM). Within one GEM all generated cDNA share a common 10x barcode. Libraries are generated and sequenced from the cDNA, and the 10x barcodes are used to associate individual reads back to the individual partitions. Incubation of the GEMs then produces barcoded, cDNA from poly-adenylated mRNA and DNA from cell surface protein Feature Barcode are generated simultaneously from the same single cell inside the GEM. After incubation, the GEMs are broken, and the pooled fractions are recovered. The cell barcoded cDNA molecules are amplified via PCR for library constructions. The amplified cDNA was then separated by SPRI size selection into cDNA fractions containing mRNA derived cDNA (>400bp) and HTO-derived cDNAs (<180bp), which were further purified by additional rounds of SPRI selection. Independent sequencing libraries were generated from the mRNA and HTO cDNA fractions, which were analyzed and quantified using TapeStation D5000 screening tapes (Agilent, Santa Clara, CA) and Qubit HS DNA quantification kit (Thermo Fisher Scientific). Libraries were pooled and sequenced together on a NovaSeq 6000 with S4 flow cell (Illumina, San Diego, CA) using paired-end, dual-index sequencing with 28 cycles for read 1, 10 cycles for i7 index, 10 cycles for i5 index, and 90 cycles for read 2.

#### Preprocessing of single-cell RNA Seq of mouse microglia

FASTQ files of single-cell sequencing libraries were processed using the “count” command of Cell Ranger (10x Genomics, version 6.0.1). Gene expression library and hashtag oligo library were processed together, and the murine transcriptome (mm10-3.1.0) was used as the reference for alignment. A mitochondrial percentage that was the higher of either 10% of reads or the 2 absolute deviations above the median for mitochondrial reads within the sample was chosen as a threshold. Cells below this threshold with greater than 500 UMIs were retained for downstream analysis. All ribosomal genes, mitochondrial genes, and pseudogenes were removed, as they interfered with the downstream differential gene expression. Hashtags were deconvoluted with demuxmix^72^. Subsequently, we checked for doublets by using a recently published tool, DoubletFinder^60^. We followed the standard workflow for DoubletFinder analysis, setting an expected doublet rate based on expected multiplet rates based on loading densities from the 10x Genomics website. The parameters we used for DoubletFinder were: 20 PCs and pN of 0.25. pK was calculated from our dataset. All doublets from demuxmix and DoubletFinder were excluded from further analysis.

To remove unwanted technical variations between experiment while preserving the variation between sex, we corrected for the batch effects separately for each sex, using scVI as described above. The scVI latent space was used for UMAP dimension reduction. Batch corrected counts were used for MELD analysis.

#### MELD analysis to quantify drug effects at the single-cell level

To quantify the heterogeneous effects of drug perturbations at a single-cell resolution, we used MELD ^23^, a graph signal processing approach that provides a continuous measure of drug effects for each cell in the transcriptomic space. The MELD score represents the relative likelihood of observing a given cell from a treatment condition of interest relative to the control. Briefly, MELD learns a low-dimensional manifold of the transcriptomic space, where cells from different treatment conditions may occupy similar or different parts of this space. The density of cells associated with each treatment condition over this manifold was then calculated (termed treatment-associated *density estimate*). To quantify the drug effect, the relative treatment-associated *likelihood* for each drug condition was calculated by normalizing the density estimate of each drug condition to that of the DMSO control. The MELD analysis was performed in python using the *meld* package (1.0.0). For the CO-iMG and iMG data, we applied MELD on the integrated space generated from the reference-query mapping procedure described above. The first 15 principal components of this integrated space were used as input to MELD. Similarly, for the mouse data, MELD was applied to the first 15 principal components of the scVI batch-corrected counts for each sex separately. To calculate the relative treatment-associated *likelihood* (herein referred as the MELD score) for each drug condition, we normalized its treatment-associated density estimates to that of the DMSO control within the same batch of the same experimental system. Key parameters *k*-nearest neighbors (*knn*) and *beta* were optimized over the ranges of 1 – 28 and 1 – 200, respectively, for each dataset using the *Benchmarker* function in *meld*. To compare the transcriptional differences among cells that were influenced by a given drug to a varying degree, we classified cells into three groups: low, medium, high, based on their levels of the MELD scores for a given drug-treated conditions. For each MELD score distribution, we pick the gating thresholds by training a 3-class gaussian mixture model using the *GaussianMixture* function of the scikit-learn package (1.2.2) in python.

#### Mitochondrial pathway analysis in human single-cell sequencing data

To delve more into the metabolic differences between compound treatment conditions in microglia isolated from the two distinct mouse models, we leveraged recent annotated murine gene sets from MitoCarta3.0 ^37^ as mentioned in **Evaluating mitotype enrichment in transcriptomic data**. Using gene sets tied to distinct mitochondrial properties, we calculated enrichment of different sets in upregulated cluster signatures using a hypergeometric test with an FDR-corrected threshold for significance at *q* = 0.01 ^66^. The results of this analysis were visualized in heatmaps where the color intensity corresponds to the -log10 p-value of the FDR q-value for enrichment of each given gene set in the differentially expressed genes associated with a given cluster.

#### Treatment of an adult amyloidosis zebrafish model with Narciclasine, Torin2 and Camptothecin

To administer the amyloid beta 42 and drugs, cerebroventricular microinjections (CVMI) into adult zebrafish brain were performed as described ^39,73^. 14 months old transgenic animals that mark microglia (*Tg (mpeg1:EGFP*); ^74^; Zebrafish International Resource Center (ZIRC)) of both sexes were used for the experiment to inject with the followings: human amyloid-beta 42 (Aβ42, 20 µM), Torin2 (1µM and 10 µM), Narciclasine (0.1 µM and 1 µM), and Camptothecin (0.1 µM and 1 µM). Total injection volume was 0.5 - 1 µl.

#### IHC staining and quantification of synaptic density, microglial density and microglial morphology in an adult amyloidosis zebrafish model

Zebrafish were euthanized at 5 days post injection; the heads were dissected and fixed overnight at 4°C using 4% paraformaldehyde (Thermofisher, Cat#: J61899-AP). After several washes the heads were incubated in 20% Sucrose (Invitrogen, Cat#: 15503-022) with 20% ethylenediaminetetraacetic acid (EDTA) (Sigma, Cat#: ED4SS) solution at 4°C for cryoprotection and decalcification, embedded in cryoprotectant sectioning resin and cryosectioned into 12μm thick sections. For immunofluorescence, the sections were dried at room temperature, followed by washing steps in PBS (Millipore Sigma, Cat#: P3813) with 0.03% Triton X-100 (Millipore Sigma, Cat#: 648466) (PBSTx). The slides were then incubated overnight with primary antibodies at 4°C. On the next day, the slides were washed 3 times with PBSTx and incubated for 2 hours in appropriate secondary antibodies at room temperature. The slides were then washed several times before mounting using aquamount (Polysciences Inc., Cat#: 18606). The following antibodies were used: chicken anti-GFP (1:1000, Invitrogen Cat#: PA1-9533), mouse anti-SV2 (1:500, DSHB Cat#: SV2, RRID:AB_2315387) and rabbit anti-L-Plastin (1:3000, ^75^, gift from Michael Redd).

Images were acquired using a Zeiss confocal LSM800 microscope in a 40X objectives with tiles/z-stack function wherever necessary. Telencephalon sections between the caudal end of the olfactory bulb and anterior commissure were used for analysis. The quantification of SV2- positive synapses was performed using 3D object counter module of ImageJ software with a same standard cut-off threshold for every images. Based on cellular morphology with respect to sphericity of DAPI+ nuclei (slender or round) and L-plastin+ microglial cell body (branched or round), three different microglial states were classified to analyze the microglial dynamics. The statistical evaluation was performed using GraphPad Prism (Version 9.5.1) for one-way ANOVA followed by a Sidak’s multiple comparison test or Dunnett’s multiple comparison test. Error bars shown are the s.e.m. and asterisks indicate significance according to: *: p<0.05, **: p<0.01, ***: p<0.001. p>0.05 is considered not significant (n.s.).

#### Structure Activity Relationship (SAR) Analysis

Compounds of interest were obtained from a wide range of reputable vendors (see Table 3) and resuspended in DMSO (Sigma-Aldrich, Cat #:472301). To keep the design of our experiment as similar as possible to the CMAP study, the target stock concentration was 10 mM. Dose titration with doses ranging from 0.01 µM to 0.1 mM was conducted to determine the highest tolerable dose for each compound. Each concentration of drug was plated in triplicate with early-passage HMC3 cells (ATCC; Cat #: CRL-3304), and cell viability was read out using using MTT assay, by incubating the cells in 0.25mg/ml MTT (invitrogen; Catalog #: M6494) in PBS (Corning, Cat #:21-040-CV) for 1hr at 37°C, following removal of MTT solution and addition of 200µl DMSO (Sigma-Aldrich, Cat #:472301) and further incubation of the cells for 15mins at 37°C before measuring the absorbance at 570nm with a Tecan Infinite® 200 PRO plate reader (Tecan; Cat#: 30050303). An optimal dose of each drug was then chosen based on cell morphology and viability. Subsequently, 0.35x10^6^ HMC3 microglial cells were seeded into a 6-well plate and incubated overnight. The next day, microglia were treated with the respective concentrations of selected drugs (see Table 3) or DMSO (Sigma-Aldrich, Cat #:472301) as control and incubated for 6hrs and 24hrs before harvest for RNA extraction. Lysis was performed in-well with buffer RLT (Qiagen; Cat #: 74136) containing 2-Mercaptoethanol (Thermo Fisher Scientific, Cat #:63689), and RNA extraction was performed with the Qiagen RNEasy mini plus kit (Qiagen; Cat #: 74136) following the manufacturer’s instructions. gDNA eliminator columns were used to remove contaminating genomic DNA. Initial RNA quality and quantity was assessed using Nanodrop (ThermoFisher Scientific) followed by cDNA preparation using the BioRad iScript cDNA Synthesis kit (BioRad; Cat #:1708891). cDNA was subsequently purified with AMPure XP beads (Thermo Fisher Scientific; Cat #: A63880) using a 1:1.8 ratio of cDNA: beads, concentration determined using Qubit HS DNA quantification kit (Thermo Fisher Scientific, Cat#: Q32851) and subsequently adjusted to 1ng/µl.

**Table 1.** Overview of compounds and concentrations used for iMG treatment.

| Compound | Vendor | Cat# | Cluster signature | Final concentration used for *in vitro* iMG experiments |
| --- | --- | --- | --- | --- |
| Torin2 | Cayman Chemical Company | 14185 | SRGAP2<sup>high</sup>/MEF2A<sup>high</sup> | 10nM |
| Narciclasine | Millipore Sigma | SML2805 | SRGAP2<sup>high</sup>/MEF2A<sup>high</sup> | 0.01μM |
| Camptothecin | EMD Millipore | 390238 | CD74<sup>high</sup>/MHC<sup>high</sup> | 0.1μM |
| Topotecan | APEX BIO | B4982 | CD74<sup>high</sup>/MHC<sup>high</sup> | 0.1μM |
| Luotonin A | Sigma-Aldrich | L4045-10MG | CD74<sup>high</sup>/MHC<sup>high</sup> | 0.1μM |

**Table 2.** Overview of compounds used for Zebrafish experiments.

| Compound | Treatment concentration | Manufacturer | Cat #: |
| --- | --- | --- | --- |
| Torin2 | 10μM and 1μM | Cayman Chemical Company | 14185 |
| Narciclasine | 1μM and 0.1μM | Millipore Sigma | SML2805 |
| Camptothecin | 1μM and 0.1 μM | EMD Millipore | 390238 |

**Table 3.**
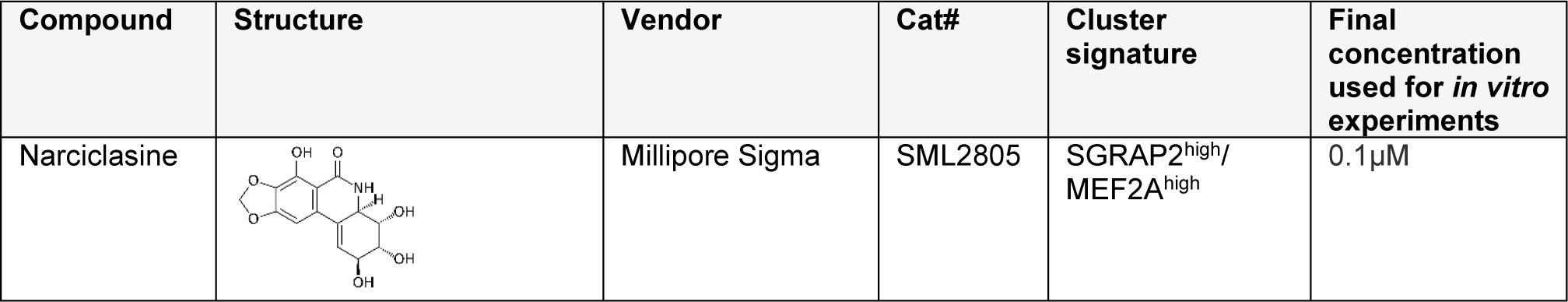
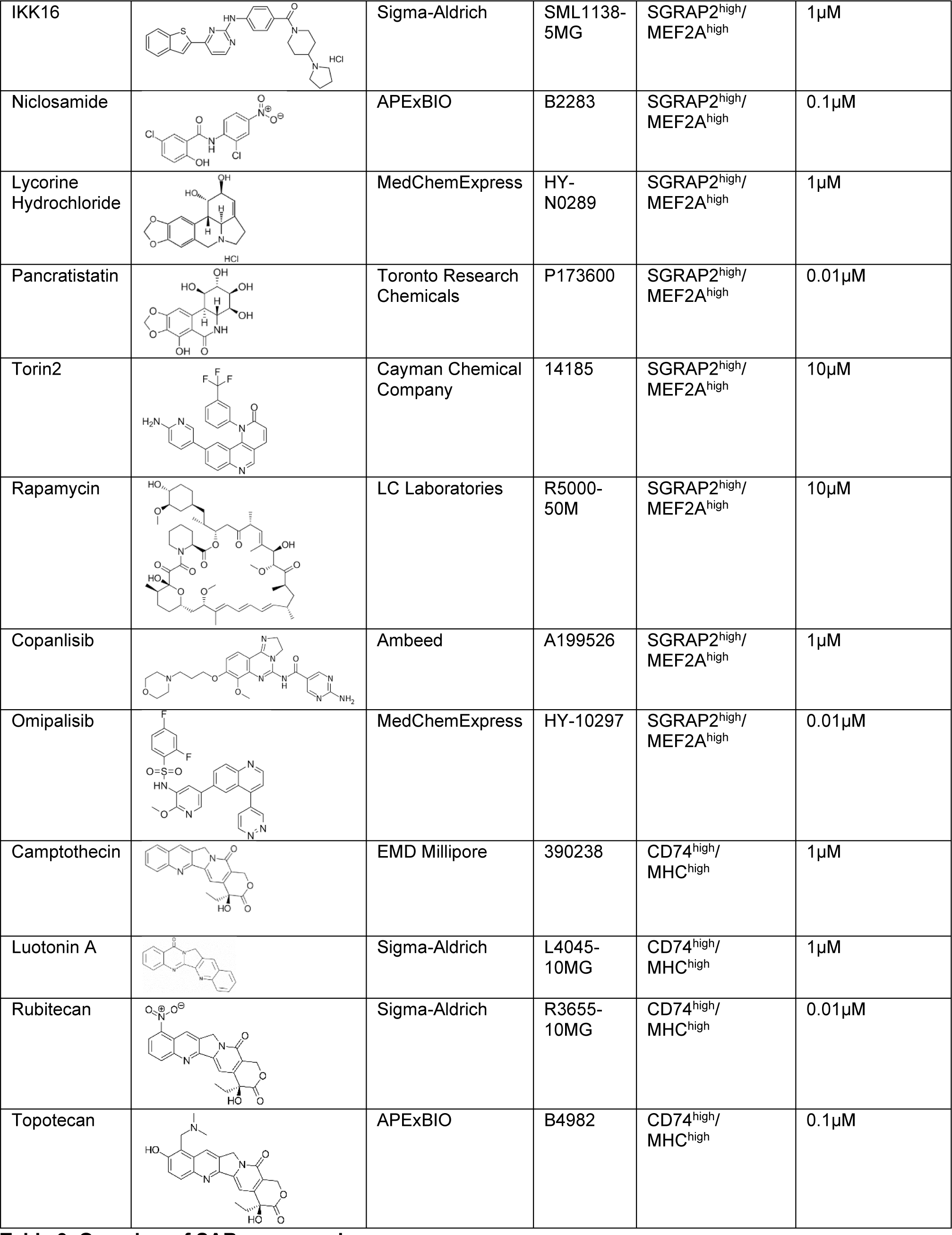
Overview of SAR compounds.

| Compound | Structure | Vendor | Cat# | Cluster signature | Final concentration used for *in vitro* experiments |
| --- | --- | --- | --- | --- | --- |
| Narciclasine |  | Millipore Sigma | SML2805 | SGRAP2<sup>high</sup>/<br>MEF2A<sup>high</sup> | 0.1µM |
| IKK16 |  | Sigma-Aldrich | SML1138-5MG | SGRAP2<sup>high</sup>/MEF2A<sup>high</sup> | 1μM |
| Niclosamide |  | APExBIO | B2283 | SGRAP2<sup>high</sup>/MEF2A<sup>high</sup> | 0.1μM |
| Lycorine Hydrochloride |  | MedChemExpress | HY-N0289 | SGRAP2<sup>high</sup>/MEF2A<sup>high</sup> | 1μM |
| Pancratistatin |  | Toronto Research Chemicals | P173600 | SGRAP2<sup>high</sup>/MEF2A<sup>high</sup> | 0.01μM |
| Torin2 |  | Cayman Chemical Company | 14185 | SGRAP2<sup>high</sup>/MEF2A<sup>high</sup> | 10μM |
| Rapamycin |  | LC Laboratories | R5000-50M | SGRAP2<sup>high</sup>/MEF2A<sup>high</sup> | 10μM |
| Copanlisib |  | Ambeed | A199526 | SGRAP2<sup>high</sup>/MEF2A<sup>high</sup> | 1μM |
| Omipalisib |  | MedChemExpress | HY-10297 | SGRAP2<sup>high</sup>/MEF2A<sup>high</sup> | 0.01μM |
| Camptothecin |  | EMD Millipore | 390238 | CD74<sup>high</sup>/MHC<sup>high</sup> | 1μM |
| Luotonin A |  | Sigma-Aldrich | L4045-10MG | CD74<sup>high</sup>/MHC<sup>high</sup> | 1μM |
| Rubitecan |  | Sigma-Aldrich | R3655-10MG | CD74<sup>high</sup>/MHC<sup>high</sup> | 0.01μM |
| Topotecan |  | APExBIO | B4982 | CD74<sup>high</sup>/MHC<sup>high</sup> | 0.1μM |

**Table 4.**
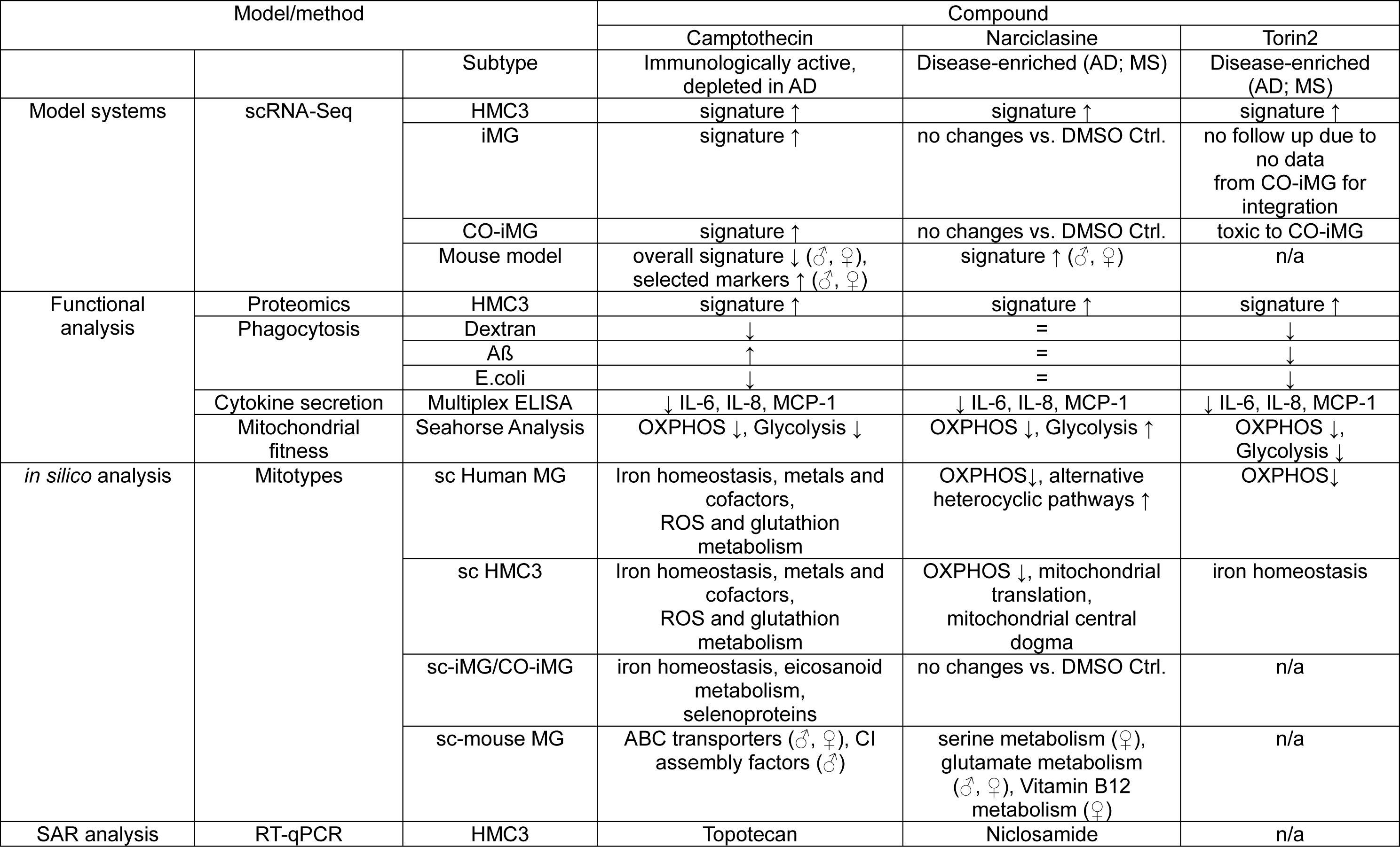

| Model/method |  |  | Compound |  |  |
| --- | --- | --- | --- | --- | --- |
|  |  |  | Camptothecin | Narciclasine | Torin2 |
|  |  | Subtype | Immunologically active, depleted in AD | Disease-enriched (AD; MS) | Disease-enriched (AD; MS) |
| Model systems | scRNA-Seq | HMC3 | signature ↑ | signature ↑ | signature ↑ |
|  |  | iMG | signature ↑ | no changes vs. DMSO Ctrl. | no follow up due to no data from CO-iMG for integration |
|  |  | CO-iMG | signature ↑ | no changes vs. DMSO Ctrl. | toxic to CO-iMG |
|  |  | Mouse model | overall signature ↓ (♂, ♀), selected markers ↑ (♂, ♀) | signature ↑ (♂, ♀) | n/a |
| Functional analysis | Proteomics | HMC3 | signature ↑ | signature ↑ | signature ↑ |
|  | Phagocytosis | Dextran | ↓ | = | ↓ |
|  |  | Aβ | ↑ | = | ↓ |
|  |  | E.coli | ↓ | = | ↓ |
|  | Cytokine secretion | Multiplex ELISA | ↓ IL-6, IL-8, MCP-1 | ↓ IL-6, IL-8, MCP-1 | ↓ IL-6, IL-8, MCP-1 |
|  | Mitochondrial fitness | Seahorse Analysis | OXPHOS ↓, Glycolysis ↓ | OXPHOS ↓, Glycolysis ↑ | OXPHOS ↓, Glycolysis ↓ |
| *in silico* analysis | Mitotypes | sc Human MG | Iron homeostasis, metals and cofactors, ROS and glutathion metabolism | OXPHOS↓, alternative heterocyclic pathways ↑ | OXPHOS↓ |
|  |  | sc HMC3 | Iron homeostasis, metals and cofactors, ROS and glutathion metabolism | OXPHOS ↓, mitochondrial translation, mitochondrial central dogma | iron homeostasis |
|  |  | sc-iMG/CO-iMG | iron homeostasis, eicosanoid metabolism, selenoproteins | no changes vs. DMSO Ctrl. | n/a |
|  |  | sc-mouse MG | ABC transporters (♂, ♀), CI assembly factors (♂) | serine metabolism (♀), glutamate metabolism (♂, ♀), Vitamin B12 metabolism (♀) | n/a |
| SAR analysis | RT-qPCR | HMC3 | Topotecan | Niclosamide | n/a |

RT- qPCR analysis was subsequently performed as described in ^12^. For cluster 1_6 analogs, the following primers were used: *SRGAP2* - fw: GTTGTGACTTAGGCTACCATGC, rev: TGCTTCGACTGTTCCAGGTTT; *MEF2A* – fw: GGTCTGCCACCTCAGAACTTT, rev:CCCTGGGTTAGTGTAGGACAA. For cluster 8_10 primers, the following primers were used: *CXCR4* – fw: ACGCCACCAACAGTCAGAG, rev: AGTCGGGAATAGTCAGCAGGA; *SRGN* – fw: GGACTACTCTGGATCAGGCTT, rev: CAAGAGACCTAAGGTTGTCATGG. *HPRT1* - fw: CCTGGCGTCGTGATTAGTGAT, rev: AGACGTTCAGTCCTGTCCATAA) was used as a housekeeping gene.

For statistical analysis, log-fold change values in comparison to DMSO-treated control samples were analyzed using one-way ANOVA followed by Dunnett’s multiple comparison test. Data was subsequently plotted in GraphPad Prism 9.2.0.

### Phagocytosis Assays

#### 1. Aβ Phagocytosis as a proxy for phagocytic behavior in a neurodegenerative context

0.1mg human beta-Amyloid (1-42), HiLyte™ Fluor 647-labeled (Anaspec Inc., Cat#: AS- 64161) was resuspended to a 4µM stock in 50µl 1% Na4OH and 3950µl Corning™ Eagle’s Minimum Essential Medium (MEM) (Corning, Cat#: 10-009-CV) and aliquots stored at -80°C until usage. For phagocytosis assay, 75K HMC3 microglia were seeded in a 24-well plate and incubated at 37°C, 5% CO2 o.n. The next day, cells were treated with Camptothecin (1µM; EMD Millipore; Cat #:390238), Narciclasine (0.1µM; Millipore Sigma; Cat #: SML2805), Torin2 (10µM; Cayman Chemical Company; Cat #: 14185) and DMSO (Sigma-Aldrich, Cat #:472301) as control and incubated for another 24hrs. Media was removed and cells were subsequently incubated in 50nM Aβ-containing complete DMEM media (10% FCS (Fisher Scientific, Cat#: 10-438-026), 1% P/S (Gibco, Cat#: 15140-122) or DMSO as a control, additionally containing either 5µM Cytochalasin D (Sigma-Aldrich, Cat#: C8273) as a negative control for phagocytosis, or DMSO as control for Cytochalasin D treatment. Cells were incubated for 1hr at 37°, 5% CO2 and subsequently washed twice with pre-warmed PBS (Corning, Cat #:21-040- CV) before harvest using Trypsin (Gen Clone; Cat #:25-510F). Cells were subsequently harvested in Flow tubes (MTC Bio, Cat#: T9005 ), centrifuged at 1500rpm for 5mins at 4°C and resuspended in 500µl Cell Staining buffer (BioLegend, Cat#: 420201) containing 1:4000 dilution of SYTOX Blue (ThermoFisher, Cat#: S34857) for labelling of live/dead cells. To assess phagocytosis, samples were subsequently assessed using Aurora 3L analyzer (Cytek Bio), followed by analysis using FlowJo_v10.8.1. Cells were gated for single cells, live cells and then Alexa647-positive cells and analyzed percentages subsequently normalized to DMSO control for each experiment to allow comparison across experiments performed on different days. For statistical analysis, log-fold change values in comparison to DMSO-treated control samples were analyzed using one-way ANOVA followed by Dunnett’s multiple comparison test. Data was subsequently plotted in GraphPad Prism 9.2.0.

#### 2. Phagocytosis of pH-rhodo Dextran as a proxy for Endocytosis

pHrhodo Green Dextran (life technologies, Cat#: P10361) was resuspended in 0.5ml deionized water (Sigma-Aldrich, Cat#: 95284) to 1mg/ml stock solution and aliquots stored at -20°C. For phagocytosis assay, 75K HMC3 microglia were seeded in a 24-well plate and incubated at 37°C, 5% CO2 o.n. The next day, cells were treated with Camptothecin (1µM; EMD Millipore; Cat #:390238), Narciclasine (0.1µM; Millipore Sigma; Cat #: SML2805), Torin2 (10µM; Cayman Chemical Company; Cat #: 14185) and DMSO (Sigma-Aldrich, Cat #:472301) as control and incubated for another 24hrs. Media was removed and cells were subsequently incubated in 100µg/ml pHrhodo Green Dextran-containing complete DMEM media (10% FCS (Fisher Scientific, Cat#: 10-438-026), 1% P/S (Gibco, Cat#: 15140-122)) or DMSO as a control, additionally containing either 5µM Cytochalasin D (Sigma-Aldrich, Cat#: C8273) as a negative control for phagocytosis, or DMSO as control for Cytochalasin D treatment. Cells were incubated for 1hr at 37°, 5% CO2 and subsequently washed twice with pre-warmed PBS (Corning, Cat #:21-040-CV) before harvest using Trypsin (Gen Clone; Cat #:25-510F) . Cells were subsequently harvested in Flow tubes (MTC Bio, Cat#: T9005 ), centrifuged at 1500rpm for 5mins at 4°C and resuspended in 500µl Cell Staining buffer (BioLegend, Cat#: 420201) containing 1:4000 dilution of SYTOX Blue (ThermoFisher, Cat#: S34857) for labeling of live/dead cells. To assess phagocytosis, samples were subsequently assessed using Aurora 3L analyzer (Cytek Bio), followed by analysis using FlowJo_v10.8.1. Cells were gated for single cells, live cells and then Alexa647-positive cells and analyzed percentages subsequently normalized to DMSO control for each experiment to allow comparison across experiments performed on different days. For statistical analysis, log-fold change values in comparison to DMSO-treated control samples were analyzed using one-way ANOVA followed by Dunnett’s multiple comparison test. Data was subsequently plotted in GraphPad Prism 9.2.0.

#### 3. Phagocytosis of pH-rhodo E.Coli as a proxy for acute neuroinflammation

pHrodo™ Green E. coli BioParticles™ Conjugate for Phagocytosis (ThermoFisher Scientific,Cat#: P35366) were resuspended in 2ml PBS (Corning, Cat #:21-040-CV) and incubated for 45mins in a sonicator bath to generate 1mg/ml stock suspension. For phagocytosis assay, 75K HMC3 microglia were seeded in a 24-well plate and incubated at 37°C, 5% CO2 o.n. The next day, cells were treated with Camptothecin (1µM; EMD Millipore; Cat #:390238), Narciclasine (0.1µM; Millipore Sigma; Cat #: SML2805), Torin2 (10µM; Cayman Chemical Company; Cat #: 14185) and DMSO (Sigma-Aldrich, Cat #:472301) as control and incubated for another 24hrs. Media was removed and cells were subsequently incubated in 160µl E.coli solution containing either 5µM Cytochalasin D (Sigma-Aldrich, Cat#: C8273) as a negative control for phagocytosis, or DMSO as control for Cytochalasin D treatment. Cells were incubated for 1hr at 37°, 5% CO2 and subsequently washed twice with pre-warmed PBS (Corning, Cat #:21-040-CV) before harvest using Trypsin (Gen Clone; Cat #:25-510F). Cells were subsequently harvested in Flow tubes (MTC Bio, Cat#: T9005 ), centrifuged at 1500rpm for 5mins at 4°C and resuspended in 500µl Cell Staining buffer (BioLegend, Cat#: 420201) containing 1:4000 dilution of SYTOX Blue (ThermoFisher, Cat#: S34857) for labelling of live/dead cells. To assess phagocytosis, samples were subsequently assessed using Aurora 3L analyzer (Cytek Bio), followed by analysis using FlowJo_v10.8.1. Cells were gated for single cells, live cells and then Alexa647-positive cells and analyzed percentages subsequently normalized to DMSO control for each experiment to allow comparison across experiments performed on different days. For statistical analysis, log-fold change values in comparison to DMSO-treated control samples were analyzed using one-way ANOVA followed by Dunnett’s multiple comparison test. Data was subsequently plotted in GraphPad Prism 9.2.0.

### Seahorse Assay to assess mitochondrial fitness

#### Oxidative Capacity and Extracellular pH

Oxygen consumption rate (OCR) and extracellular acidification rate (ECAR; pH change) were measured over a confluent cell monolayer of HMC3 microglia using the XFe96 Seahorse extracellular flux analyzer (Agilent Technologies). First, 10K HMC3 microglia per well were seeded in 80µl of a Agilent Seahorse XF96 Cell Culture Microplate (Agilent Technologies; Cat #: 101085-004) and incubated overnight. The next day, cells were treated with respective compounds, including Camptothecin (1µM; EMD Millipore; Cat #:390238), Narciclasine (0.1µM; Millipore Sigma; Cat #: SML2805), Torin2 (10µM; Cayman Chemical Company; Cat #: 14185) or DMSO (Sigma-Aldrich, Cat #:472301) as control in a total volume of 80µl/ well and incubated for 24hrs. Subsequently, OCR and ECAR were measured as follows: cell culture media was removed and 80µl of XF Media (Agilent Technologies; Cat #: 103334-100) containing no pH buffers and supplemented with 5.5 mM glucose , 1 mM pyruvate (Sigma-Aldrich; Cat #: Sigma-Aldrich #P9767), 1 mM glutamine (fisher scientific; Cat #: G7513), 50 µg/ml Uridine (Sigma-Aldrich, Cat#: U6381), 10 µM palmitate (Sigma-Aldrich, Cat#: P9767) conjugated to 1.7 µM BSA (pluriSelect; Cat#: 60-00020-10) was added to the cells. The plate was then incubated in a non-CO2 incubator for one hour to equilibrate temperature and atmospheric gases. The Seahorse instrument was programmed to assess various respiratory states using the manufacturer’s protocol of sequential substrate addition and measurements (4x, 3 measurements over 18 mins). Basal respiration, ATP turnover, proton leak, coupling efficiency, maximum respiration rate, respiratory control ratio, spare respiratory capacity and non-mitochondrial respiration were all determined by the sequential additions of the ATP synthase inhibitor oligomycin (final concentration: 1 µM; Sigma-Aldrich Cat#:75351), the protonophoric uncoupler FCCP (final concentration: 0.45 µM; Selleckchem, Cat #: S8276), and the electron transport chain Complex I and III inhibitors, rotenone (final concentration: 1 µM ; Sigma-Aldrich, Cat #: R8875) and antimycin A (final concentration: 1 µM ; Sigma-Aldrich, Cat #: A8674) respectively as well as the staining dye Hoechst (1:200 dilution; ThermoFisher Scientific #62249). The optimal concentration for the uncoupler FCCP yielding maximal uncoupled respiration was previously determined based on a titration performed on HMC3 microglia (**see Supplemental Figure 4**).

After the seahorse run, nuclei were counted using the cytation1 Cell Imager (BioTek). Both OCR and ECAR were normalized relative cell counts on a per-well basis. Bioenergetic metrics were calculated using the P/O ratios of OxPhos and glycolysis as described in ^32^. These calculations assume that the pyruvate was derived entirely from glucose via glycolysis. The code and raw data are available as detailed in the Data Availability statement.

#### Cytokine Multiplex Assay

10K HMC3 cells from three different passages (P12, P19, P21) were seeded in 96-well plates containing 200µl of complete Eagle’s Minimum Essential Medium (EMEM; ATCC, Cat#: 30- 2003) and incubated o.n. at 37°C, 5% CO2. The next day were treated with compounds using the following concentrations (6 wells per condition): DMSO (1:1000), Narciclasine (0.1µM), Torin2 (10µM), Camptothecin (1µM), Topotecan (1µM), Rubitecan (0.01µM), Luotonin A (1µM) and incubated for 24hrs. The next day, 2 wells/ condition were stimulated with TNF-a (0.3 µg/mL, Peprotech, Cat#: 300-02), 2 wells/ condition were stimulated with IFN-y (0.3 µg/mL, Peprotech, Cat#: 300-01A) and as a control 2 wells/condition were stimulated with H2O as control, each dissolved in a fresh dilution of the respective drug treatment in a final volume of 200µl. For subsequent cytokine expression analysis, cell culture supernatant was harvested after 12hrs and 24hrs and stored at -20°C until further analysis.

For cytokine multiplex assay, 100µl/sample were sent to Eve Technologies (Calgary, Alberta) and analyzed using Human Cytokine Proinflammatory Focused 15-Plex Discovery Assay® Array (HDF15). Data was subsequently plotted in GraphPad Prism 9.2.0. For statistical analysis, one-way ANOVA followed by Tukey’s multiple comparisons test with a single pooled variance was performed using GraphPad Prism 9.2.0 software.

