## Supplementary Information for "A pharmacological toolkit for human microglia identifies Topoisomerase I inhibitors as immunomodulators for Alzheimer’s disease"

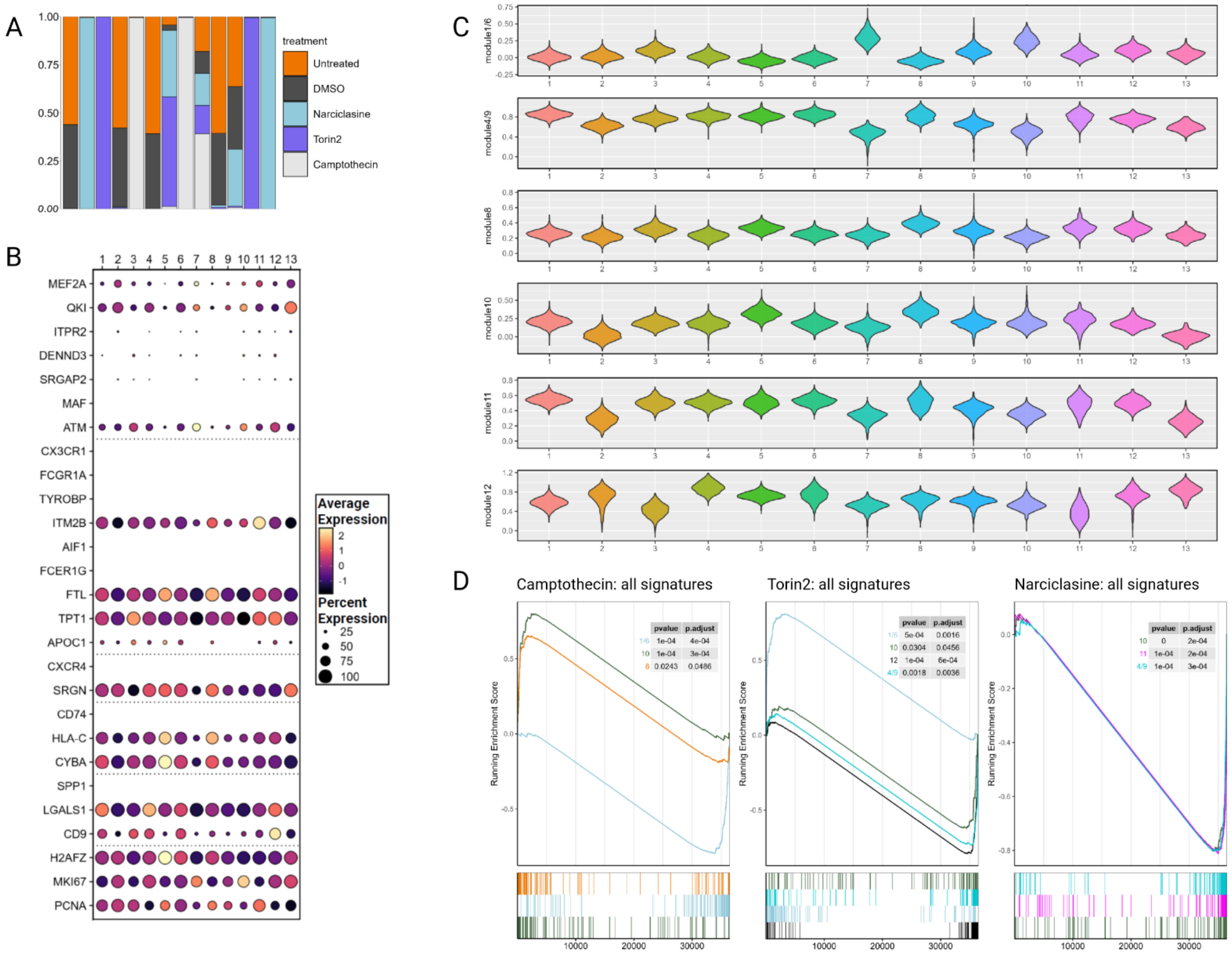


**Supplementary Figure 1. A.** Bar plots showing the relative representation of different clusters across different treatment conditions, with each bar summing to 100%. **B. Microglial markers projected on HMC3 scRNA-seq clusters.** Each column represents data from a given cluster, each row represents the expression level of each gene. Circle size indicates the percentage of cells per each cluster expressing the gene. The circle color represents z-scored gene expression **C. Expression** levels of different expression modules of human microglial subpopulations across compound-treated HMC3 microglial subclusters. Violin plots depict the per-cell module scores for CD74$^{high}$/MHC$^{high}$ and SRGAP2$^{high}$/MEF2A$^{high}$ microglial subclusters as well as modules 4/9, 8, 11, 12 derived from [1], grouped by de novo defined HMC3 microglial subclusters 1-13. **D. Gene set enrichment analysis for different module expression levels in compound-treated HMC3 microglia.** Each plot shows gene set enrichment scores for distinct modules relevant to each drug treatment – Camptothecin (modules 1/6, 8, 10), Torin2 (modules 1/6, 4/9, 10, 12), Narciclasine (modules 4/9, 10, 11).

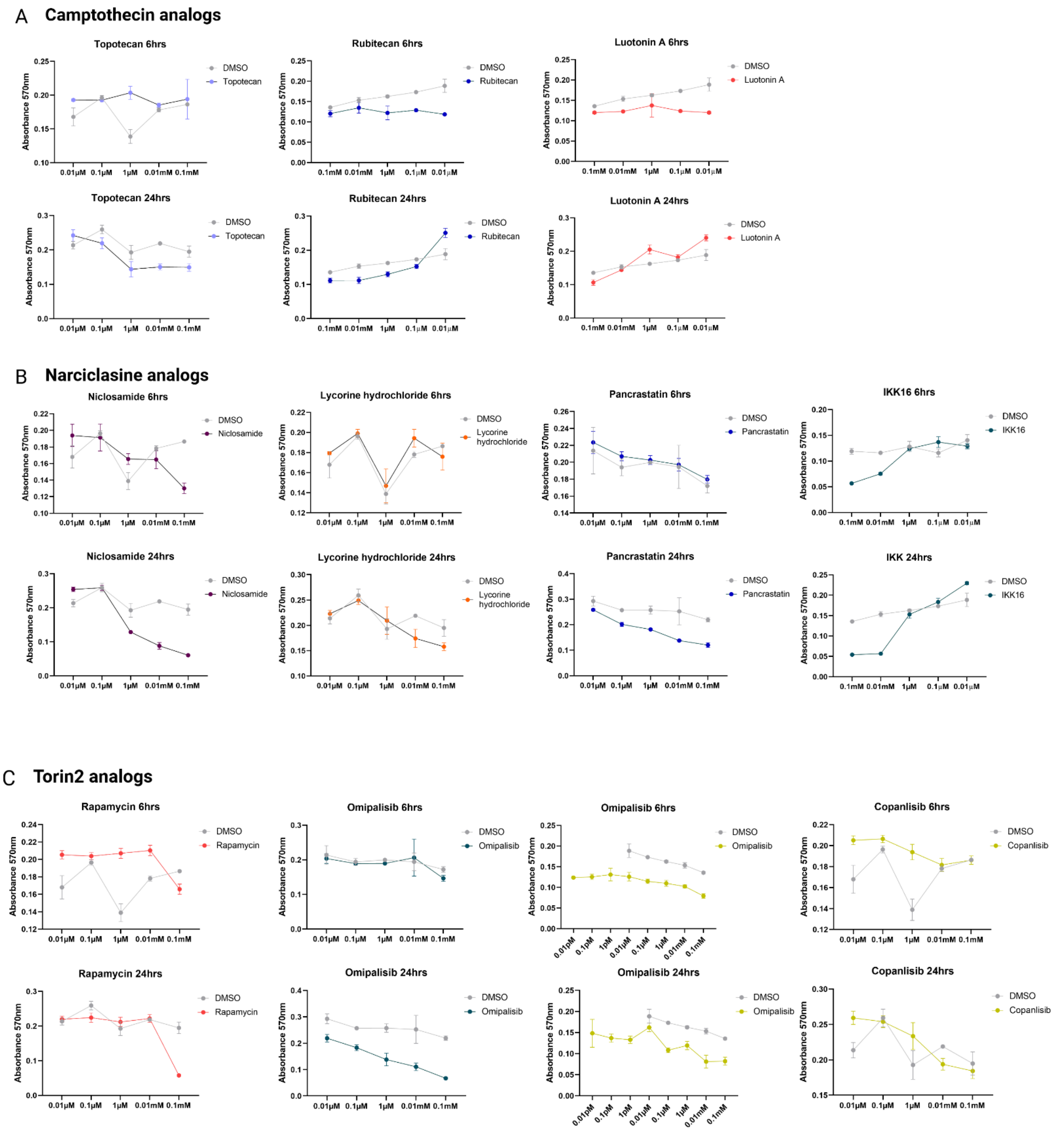
A Camptothecin analogs
Topotecan 6hrs
Rubitecan 6hrs
Luotonin A 6hrs
Topotecan 24hrs
Rubitecan 24hrs
Luotonin A 24hrs
DMSO
Topotecan
Rubitecan
Luotonin A
B Narciclasine analogs
Niclosamide 6hrs
Lycorine hydrochloride 6hrs
Pancrastatin 6hrs
IKK16 6hrs
Niclosamide 24hrs
Lycorine hydrochloride 24hrs
Pancrastatin 24hrs
IKK 24hrs
Niclosamide
Lycorine hydrochloride
Pancrastatin
IKK16
C Torin2 analogs
Rapamycin 6hrs
Omipalisib 6hrs
Copanlisib 6hrs
Rapamycin 24hrs
Omipalisib 24hrs
Copanlisib 24hrs
Rapamycin
Omipalisib
Copanlisib
Absorbance 570nm
0.01μM 0.1μM 1μM 0.01mM 0.1mM
0.1mM 0.01mM 1μM 0.1μM 0.01μM

D

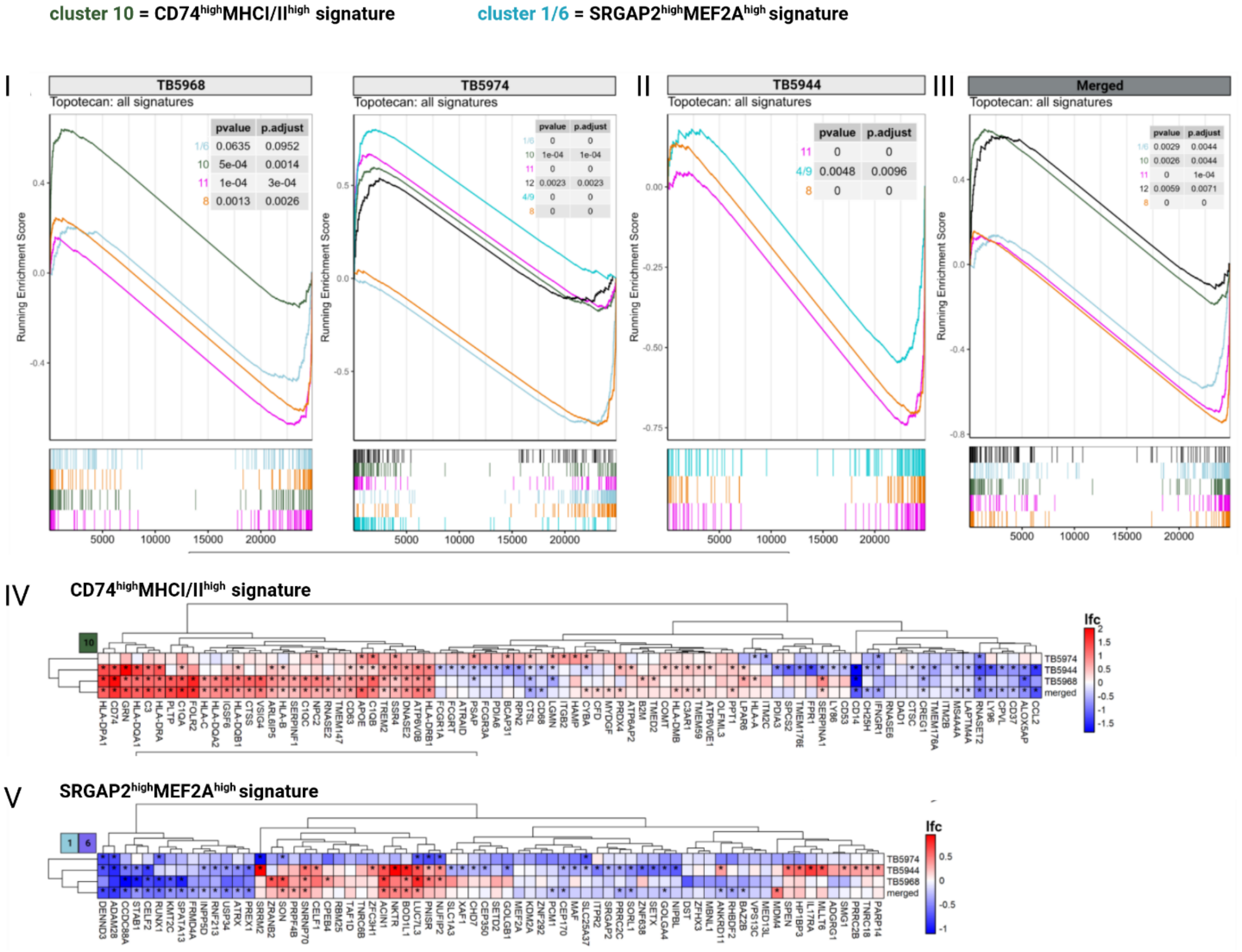


**Supplementary Figure 2. A. Titration experiments of SAR (structure activity relationship analysis).** Each compound was titrated on HMC3 microglia covering a range from 0.1mM to 0.01µM (0.1mM, 0.01mM, 1µM, 0.1µM, 0.01µM). Following 6hrs and 24hrs treatment, cell viability was assessed using MTT viability assay. **A.** MTT viability assay for Camptothecin analogs (Topotecan, Rubitecan, Luotonin A) 6hrs and 24hrs following treatment, in comparison to DMSO control. Data points/concentration are derived from treatment in triplicates as mean ± SD. Compound-treatment is depicted in color, DMSO controls are visualized in grey. **B.** MTT viability assay for Narciclasine analogs (Niclosamide, Lycorine Hydrochloride, Pancrastatin, IKK16) 6hrs and 24hrs following treatment in comparison to DMSO control. Data points/concentration are derived from treatment in triplicates as mean ± SD. Compound-treatment is depicted in color, DMSO controls are visualized in grey. **C.** MTT viability assay for Torin2 analogs (Rapamycin, Omipalisib, Copanlisib) 6hrs and 24hrs following treatment, in comparison to DMSO control. For Omipalisib, additional titrations between 0.01pM – 1pM were performed (0.01pM, 0.1pM, 1pM). Data points/concentration are derived from treatment in triplicates as mean ± SD. Compound-treatment is depicted in color, DMSO controls are visualized in grey. **D. Topotecan treatment induces CD74$^{high}$/MHC$^{high}$ signature and reduces SRGAP2$^{high}$/MEF2A$^{high}$ signature on a per-sample basis in myeloid cells isolated from primary human glioblastoma slice cultures. I. GSEA demonstrates upregulation of the CD74$^{high}$/MHC$^{high}$ signature in patient-derived samples TB5968 and TB5974.** GSEA was conducted evaluating the enrichment of the top 100 genes per cluster or cluster grouping in the ordered list of genes detected in our dataset. Genes were ordered based on average log-fold change between cells from the Topotecan treatment condition versus control. The running enrichment score as well as the position of each gene in the ranked list is shown. **II. GSEA does not demonstrate upregulation of the cluster 10 signature in TB5944. III. GSEA on merged data demonstrates consistent upregulation of the CD74$^{high}$/MHC$^{high}$ signature and downregulation of the SRGAP2$^{high}$/MEF2A$^{high}$ signature.** The three prior samples were merged for this analysis. **IV. Topotecan consistently upregulates complement/MHC genes across samples.** Heatmap depicts log-fold change of Topotecan vs. control samples across three different samples and in the merged object. Rows are individual samples, columns represent genes. Clustering was performed using hierarchical clustering by maximum linkage.

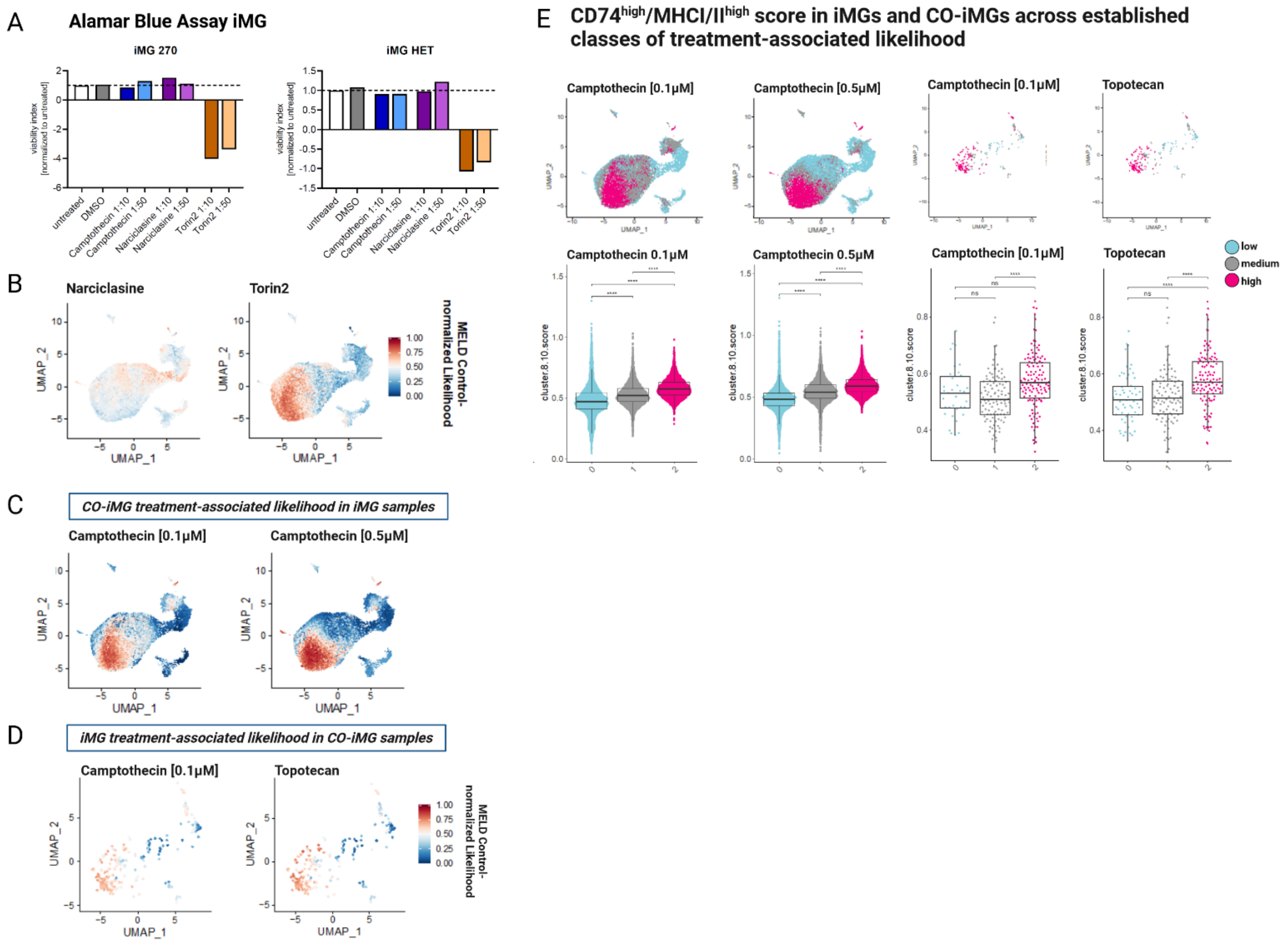


**Supplementary Figure 3. A. Alamar blue assay assessing viability of iMGs after 24hrs of treatment with selected compounds** or DMSO-treated and untreated cells as control (Camptothecin (0.1µM; 0.02µM); Narciclasine (10nM, 2nM), Torin 2 (1µM, 0.2µM)). Graphs depict assessed viability index for 1n of two distinct iPSC-derived microglia (iMG) lines (270 and HET **B. Individual iMGs colored by treatment-associated likelihood in the joint UMAP.** Cells are colored by the control-normalized treatment-associated likelihood calculated by MELD [2] for Narciclasine- and Torin2-treated iMGs. For a given treatment condition, the relative treatment likelihood represents the likelihood of observing a cell from the given treatment condition (red) relative to the DMSO control (blue). **C. Individual iMGs and CO-iMGs colored by treatment-associated likelihood in the joint UMAP.** Cells (including iMGs) are colored by the control-normalized treatment-associated likelihood calculated by MELD for the CO-iMGs treatment conditions with 0.1µM Camptothecin and 0.5 µM Camptothecin. **D. Individual iMGs and CO-iMGs colored by treatment-associated likelihood in the joint UMAP.** CO-iMGs are shown and colored the control-normalized treatment-associated likelihood calculated by MELD for the iMGs treatment conditions with 0.1µM Camptothecin and Topotecan. **E. CD74$^{high}$/MHC$^{high}$ score in iMGs and CO-iMGs across different levels of treatment-associated likelihood.** Cells are colored by three different levels (low, medium, high) of treatment-associated likelihood for each drug-treatment condition. The first two plots from the left show the iMGs cells gated by the treatment-associated likelihood inferred from Camptothecin-treated CO-iMG data. The second two plots show only CO-iMGs in the same UMAP space, where each CO-iMGs is colored by the Camptothecin and Topotecan treatment-associated likelihood calculated from the iMG data. Bottom row - single-cell distributions of the Cluster 8/10 gene module score [1] across the three levels of treatment-associated likelihood inferred from the indicated treatment conditions. Each boxplot highlights the median, lower and upper quartiles. Whiskers in boxplots indicate 1.5 times interquartile ranges.

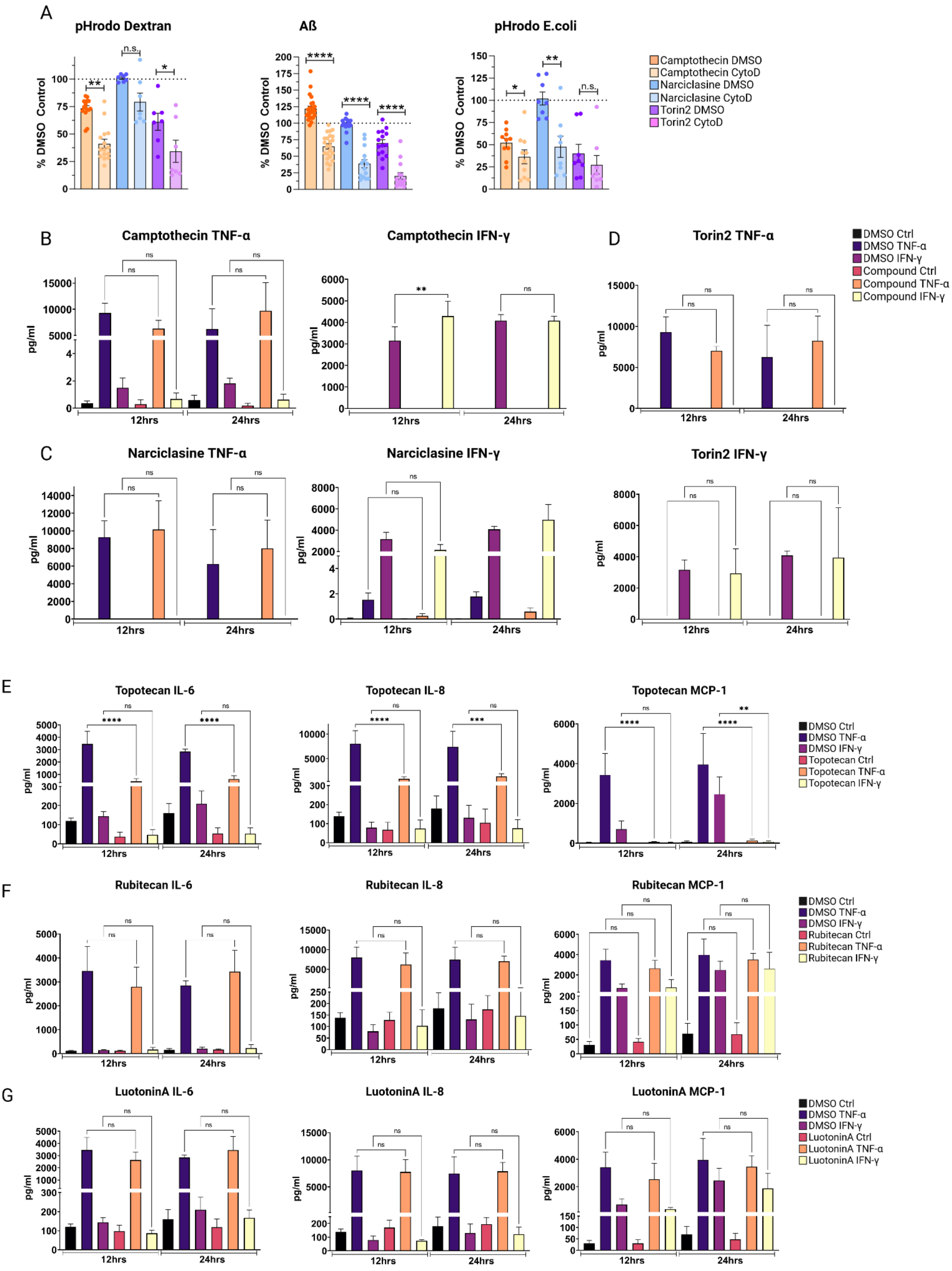
A
pHrodo Dextran
Aß
pHrodo E.coli
% DMSO Control
Camptothecin DMSO
Camptothecin CytoD
Narciclasine DMSO
Narciclasine CytoD
Torin2 DMSO
Torin2 CytoD
B
Camptothecin TNF-α
Camptothecin IFN-γ
D
Torin2 TNF-α
DMSO Ctrl
DMSO TNF-α
DMSO IFN-γ
Compound Ctrl
Compound TNF-α
Compound IFN-γ
C
Narciclasine TNF-α
Narciclasine IFN-γ
Torin2 IFN-γ
pg/ml
12hrs
24hrs
E
Topotecan IL-6
Topotecan IL-8
Topotecan MCP-1
Topotecan Ctrl
Topotecan TNF-α
Topotecan IFN-γ
F
Rubitecan IL-6
Rubitecan IL-8
Rubitecan MCP-1
Rubitecan Ctrl
Rubitecan TNF-α
Rubitecan IFN-γ
G
LuotoninA IL-6
LuotoninA IL-8
LuotoninA MCP-1
LuotoninA Ctrl
LuotoninA TNF-α
LuotoninA IFN-γ

H Representative Seahorse Traces

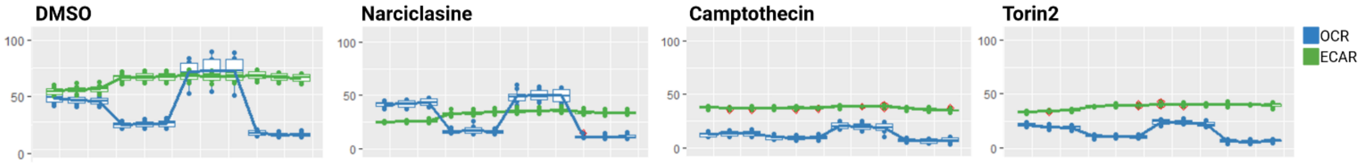


**Supplementary Figure 4. A. Complete figures of endocytosis and phagocytosis assays following compound-treatment (Camptothecin, Narciclasine, Torin2, DMSO as control) using different substrates.** For all three assays, HMC3 microglia were pretreated with respective compounds (Camptothecin, Narciclasine, Torin2) or DMSO as control for 24hrs, subsequently exposed to pHrodo-labeled Dextran, Alexa 647-labelled Aβ1-42 or pHrodo-labeled E.coli either containing either 5μM Cytochalasin D or DMSO as control for Cytochalasin D treatment. After 1hr the uptake of the respective substrate was assessed using flow cytometry. Individual experiments are depicted as individual dots in the bar graphs depicting mean ± SEM (Camptothecin – orange; Narciclasine – blue; Torin2 - purple). Phagocytosis was normalized to % DMSO control and for statistical analysis, log-fold change values of compound- and Cytochalasin D -treated cells in comparison to compound-treated samples was analyzed using Mann-Whitney U test. *p.adj ≤ 0.05; **p.adj ≤ 0.01; ***p.adj ≤ 0.001; ****p.adj ≤ 0.0001. **B-D.** Effect of **Narciclasine, Torin2 and Camptothecin on the secretion of pro-inflammatory cytokines TNF-α and IFN-y.** HMC3 microglia were pre-treated with Narciclasine, Torin2 and Camptothecin for 24hrs, and subsequently stimulated with either TNF-a (0.3 μg/mL), IFN-y (0.3 μg/mL) or $H_2O$ as control for 12 or 24hrs. Supernatant was collected and pro-inflammatory cytokine secretion assessed using a human pro-inflammatory cytokine discovery assay. Bar graphs depict measured amount of cytokines TNF-α and IFN-y (mean ± SEM) in pg/ml for DMSO control-treated samples (black, dark purple, light purple) or compound-treated samples (red, orange, yellow). For statistical analysis, one-way ANOVA followed by Tukey's multiple comparisons test with a single pooled variance was performed. *p.adj ≤ 0.05; **p.adj ≤ 0.01; ***p.adj ≤ 0.001; ****p.adj ≤ 0.0001. **E-G.** Effect of **SAR compounds (Topotecan, Rubitecan, Luotonin A) on the secretion of pro-inflammatory cytokines.** HMC3 microglia were pre-treated with Topotecan, Rubitecan or Luotonin A for 24hrs, and subsequently stimulated with either TNF-a (0.3 μg/mL), IFN-y (0.3 μg/mL) or $H_2O$ as control for 12 or 24hrs. Supernatant was collected and pro-inflammatory cytokine secretion assessed using a human pro-inflammatory cytokine discovery assay. Bar graphs depict measured amount of IL-6, IL-8, MCP-1 (mean ± SEM) in pg/ml for DMSO control-treated samples (black, dark purple, light purple) or compound-treated samples (red, orange, yellow). For statistical analysis, one-way ANOVA followed by Tukey's multiple comparisons test with a single pooled variance was performed. *p.adj ≤ 0.05; **p.adj ≤ 0.01; ***p.adj ≤ 0.001; ****p.adj ≤ 0.0001. **H.** Representative seahorse assay traces depicting oxygen consumption rate (OCR, blue) and extracellular acidification rate (ECAR, green).

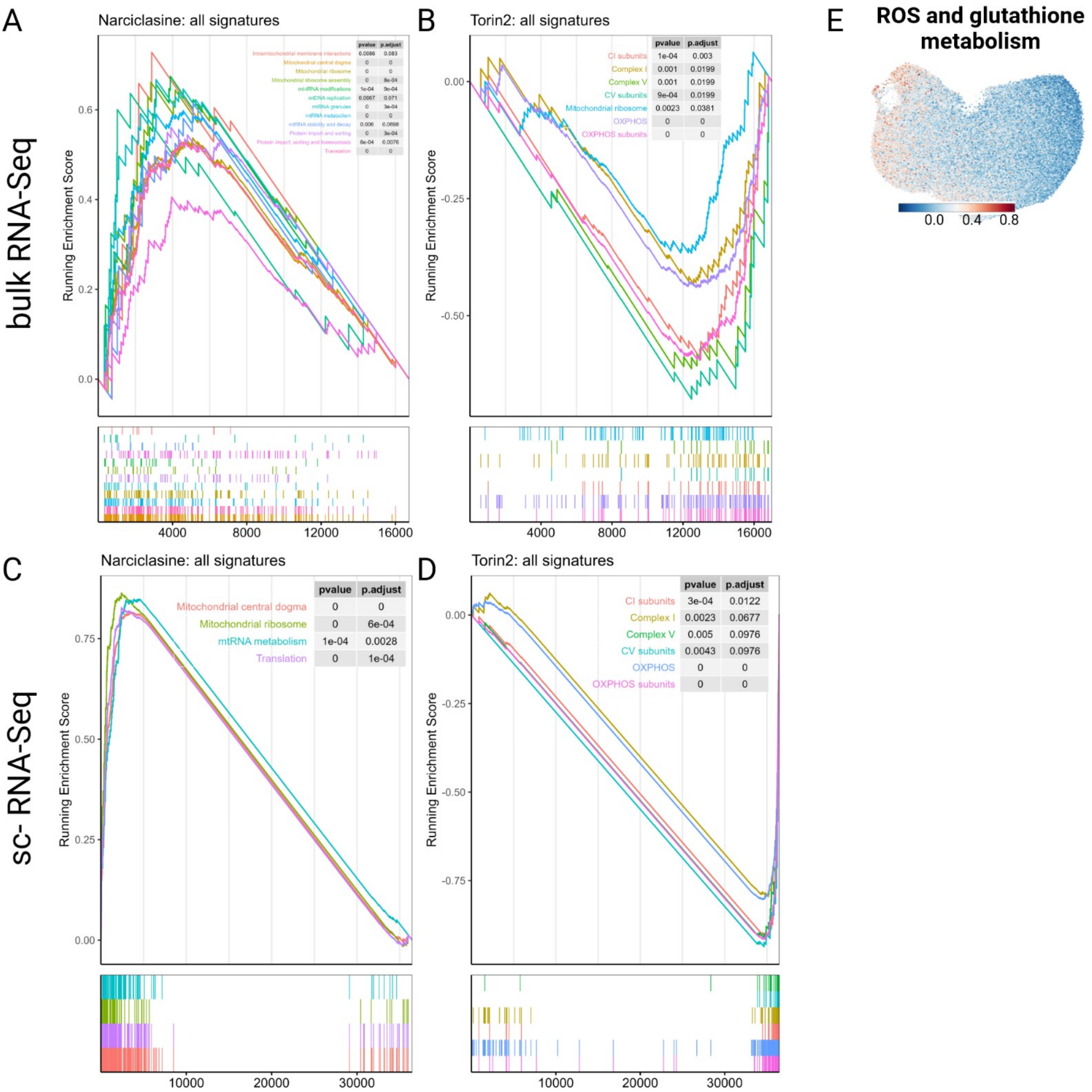
A
Narciclasine: all signatures
bulk RNA-Seq
Running Enrichment Score
pvalue p.adjust
Intramitochondrial membrane interactions 0.0086 0.083
Mitochondrial central dogma 0 0
Mitochondrial ribosome 0 0
Mitochondrial ribosome assembly 0 8e-04
mt-tRNA modifications 1e-04 9e-04
mtDNA replication 0.0067 0.071
mtRNA granules 0 3e-04
mtRNA metabolism 0 0
mtRNA stability and decay 0.006 0.0698
Protein import and sorting 0 3e-04
Protein import, sorting and homeostasis 6e-04 0.0076
Translation 0 0
0.6
0.4
0.2
0.0
4000
8000
12000
16000
B
Torin2: all signatures
Running Enrichment Score
pvalue p.adjust
CI subunits 1e-04 0.003
Complex I 0.001 0.0199
Complex V 0.001 0.0199
CV subunits 9e-04 0.0199
Mitochondrial ribosome 0.0023 0.0381
OXPHOS 0 0
OXPHOS subunits 0 0
0.00
-0.25
-0.50
4000
8000
12000
16000
C
Narciclasine: all signatures
sc- RNA-Seq
Running Enrichment Score
pvalue p.adjust
Mitochondrial central dogma 0 0
Mitochondrial ribosome 0 6e-04
mtRNA metabolism 1e-04 0.0028
Translation 0 1e-04
0.75
0.50
0.25
0.00
10000
20000
30000
D
Torin2: all signatures
Running Enrichment Score
pvalue p.adjust
CI subunits 3e-04 0.0122
Complex I 0.0023 0.0677
Complex V 0.005 0.0976
CV subunits 0.0043 0.0976
OXPHOS 0 0
OXPHOS subunits 0 0
0.00
-0.25
-0.50
-0.75
10000
20000
30000
E
ROS and glutathione metabolism
0.0
0.4
0.8

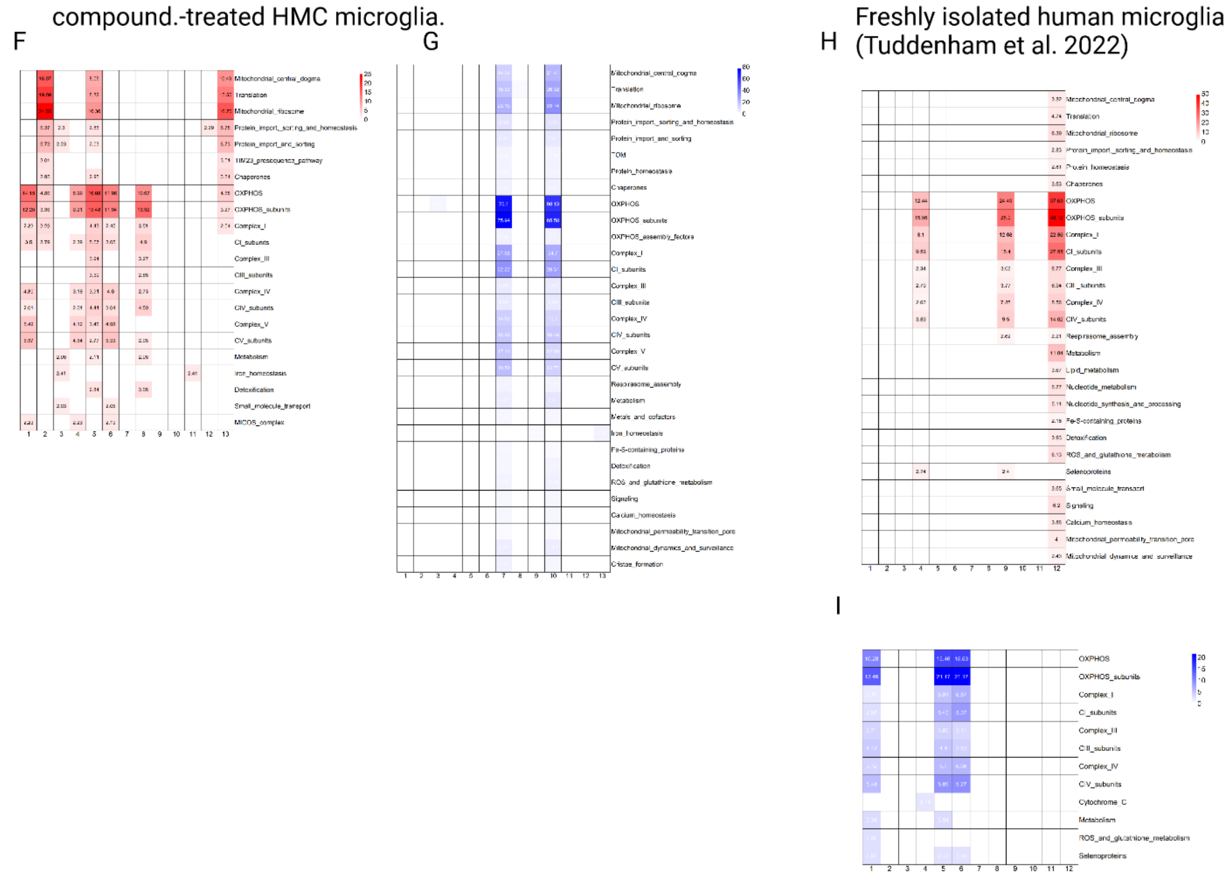


**Supplementary Figure 5. Mitochondrial phenotyping of compound-treated HMC3s. A. Narciclasine upregulates mitochondrial translation and activity at the bulk RNA level.** Enrichment of modules of genes associated with different aspects of mitochondrial activity was performed using GSEA in the ordered list of genes detected in our dataset, ordered based on average log-fold change between cells from the treatment condition vs. control. The running enrichment score as well as the position of each gene in the ranked list is shown. **B. Torin-2 downregulates oxidative phosphorylation at the bulk RNA level. C. Narciclasine upregulates mitochondrial translation at the single-cell level.** Here, all single cells from each condition were aggregated into treatment or control (both DMSO and untreated) and the mean log fold change was calculated. **D. Torin-2 also downregulates oxidative phosphorylation at the single cell level**. **E.** Selected results of GSEA for specific mitotypes in human microglial subtypes are shown. Legend depicts module score, showing average expression for module genes vs. a background of similarly expressed control genes. **F. Mitochondrial phenotyping identifies clusters enriched in different aspects of respiratory metabolism (compound-treated HMC3 microglia).** Enrichment of modules of genes associated with different aspects of mitochondrial activity was performed in upregulated gene lists associated with each cluster (compound-treated HMC3 microglia) using the hypergeometric test with Benjamini-Hochberg correction. Bars are colored if they have an FDR < 0.01. **G. Mitochondrial phenotyping identifies clusters downregulated in different aspects of respiratory metabolism (compound-treated HMC3 microglia).** Enrichment of modules of genes associated with different aspects of mitochondrial activity was performed in downregulated gene lists associated with each cluster (compound-treated HMC3 microglia) using the hypergeometric test with Benjamini-Hochberg correction. Bars are colored if they have an FDR < 0.01. **H. Mitochondrial phenotyping identifies clusters enriched in different aspects of respiratory metabolism (freshly isolated human microglia; [1]).** Enrichment of modules of genes associated with different aspects of mitochondrial activity was performed in upregulated gene lists associated with each cluster (freshly isolated human microglia;[1]) using the hypergeometric test with Benjamini-Hochberg correction. Bars are colored if they have an FDR < 0.01. **I. Mitochondrial phenotyping identifies clusters downregulated in different aspects of respiratory metabolism.** Enrichment of modules of genes associated with different aspects of mitochondrial activity was performed in downregulated gene lists associated with each cluster (freshly isolated human microglia; [1]) using the hypergeometric test with Benjamini-Hochberg correction. Bars are colored if they have an FDR < 0.01.

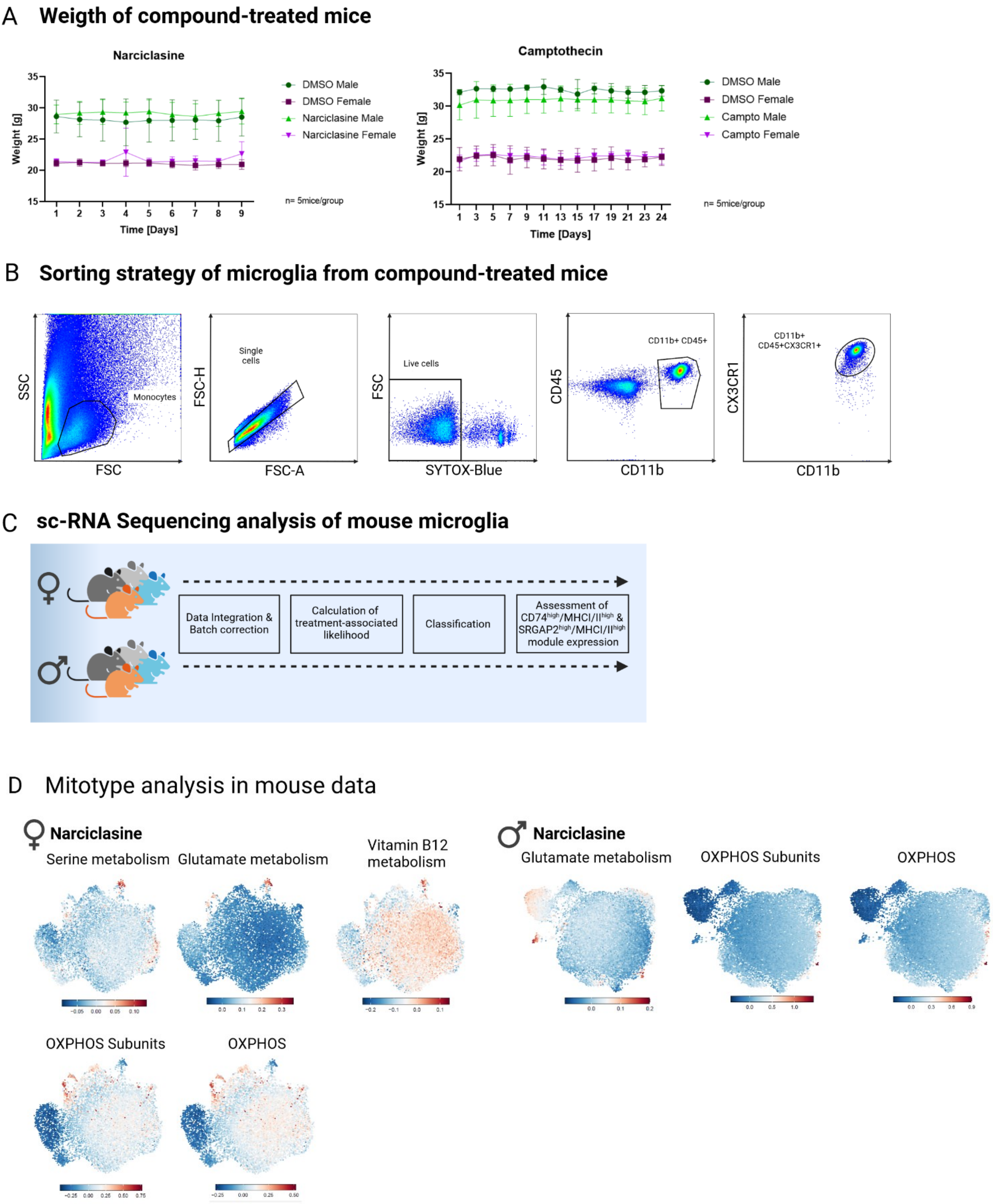
A Weigth of compound-treated mice
Narciclasine
Weight [g]
Time [Days]
DMSO Male
DMSO Female
Narciclasine Male
Narciclasine Female
n= 5mice/group
Camptothecin
Weight [g]
Time [Days]
DMSO Male
DMSO Female
Campto Male
Campto Female
n= 5mice/group
B Sorting strategy of microglia from compound-treated mice
SSC
FSC
Monocytes
FSC-H
FSC-A
Single cells
FSC
SYTOX-Blue
Live cells
CD45
CD11b
CD11b+ CD45+
CX3CR1
CD11b
CD11b+ CD45+CX3CR1+
C sc-RNA Sequencing analysis of mouse microglia
Data Integration & Batch correction
Calculation of treatment-associated likelihood
Classification
Assessment of CD74^high/MHCI/II^high & SRGAP2^high/MHCI/II^high module expression
D Mitotype analysis in mouse data
Narciclasine
Serine metabolism
Glutamate metabolism
Vitamin B12 metabolism
OXPHOS Subunits
OXPHOS
Narciclasine
Glutamate metabolism
OXPHOS Subunits
OXPHOS

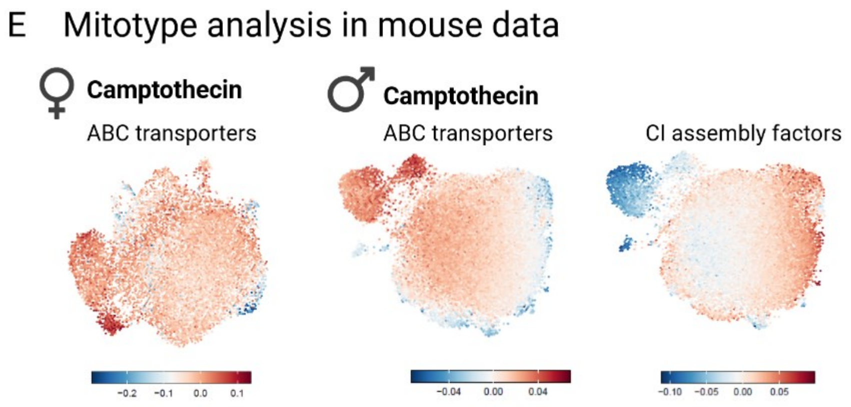


**Supplementary Figure 6.A. Weight of compound-treated mice over time.** Graphs depict weight of mice (g) over treatment time (days) for Narciclasine-treated mice for 8 days of treatment and on day 9 of experimental stop, for Camptothecin-treated mice for 23 days and on day 24 of experimental stop. Males are depicted in green (DMSO control, dark green; treatment in light green), females are depicted in purple (DMSO control, dark purple; treatment in light purple). n=5 mice/treatment group. **B. Sorting strategy of microglia from compound-treated mice.** Following FSC and SSC-based gating, single cell and live cell gating based on Sytox-Blue staining, microglia were gated as CD11b+CD45+ cells and further gated for CX3CR1+ expression. Cells with CD11b+CD45+CX3CR1+ identity were then sorted for subsequent sc-RNA sequencing. **C. Analysis strategy of sc-RNA Seq data from compound-treated mice.** For analysis, female and male microglia from all treatment groups were analyzed jointly but separated by sex. Female or male microglia from all treatment conditions were integrated, followed by batch correction and treatment-associated likelihood was calculated using the MELD algorithm [2]. Based on treatment-associated likelihood, cells were then classified into low, medium and high classes and the expression of $CD74^{high}$/$MHC^{high}$ and $SRGAP^{high}$/$MEF2A^{high}$ signatures was assessed. **D. Mitotype analysis in microglia isolated from male and female mice treated with Narciclasine.** Enrichment of modules of genes associated with different aspects of mitochondrial activity was performed using GSEA in the ordered list of genes detected in our dataset. Genes in our data were ordered based on average log-fold change between cells from the treatment condition versus control. **E. Mitotype analysis in microglia isolated from male and female Camptothecin- treated mice.** Enrichment of modules of genes associated with different aspects of mitochondrial activity was performed using GSEA in the ordered list of genes detected in our dataset. Genes in our data were ordered based on average log-fold change between cells from the treatment condition versus control.

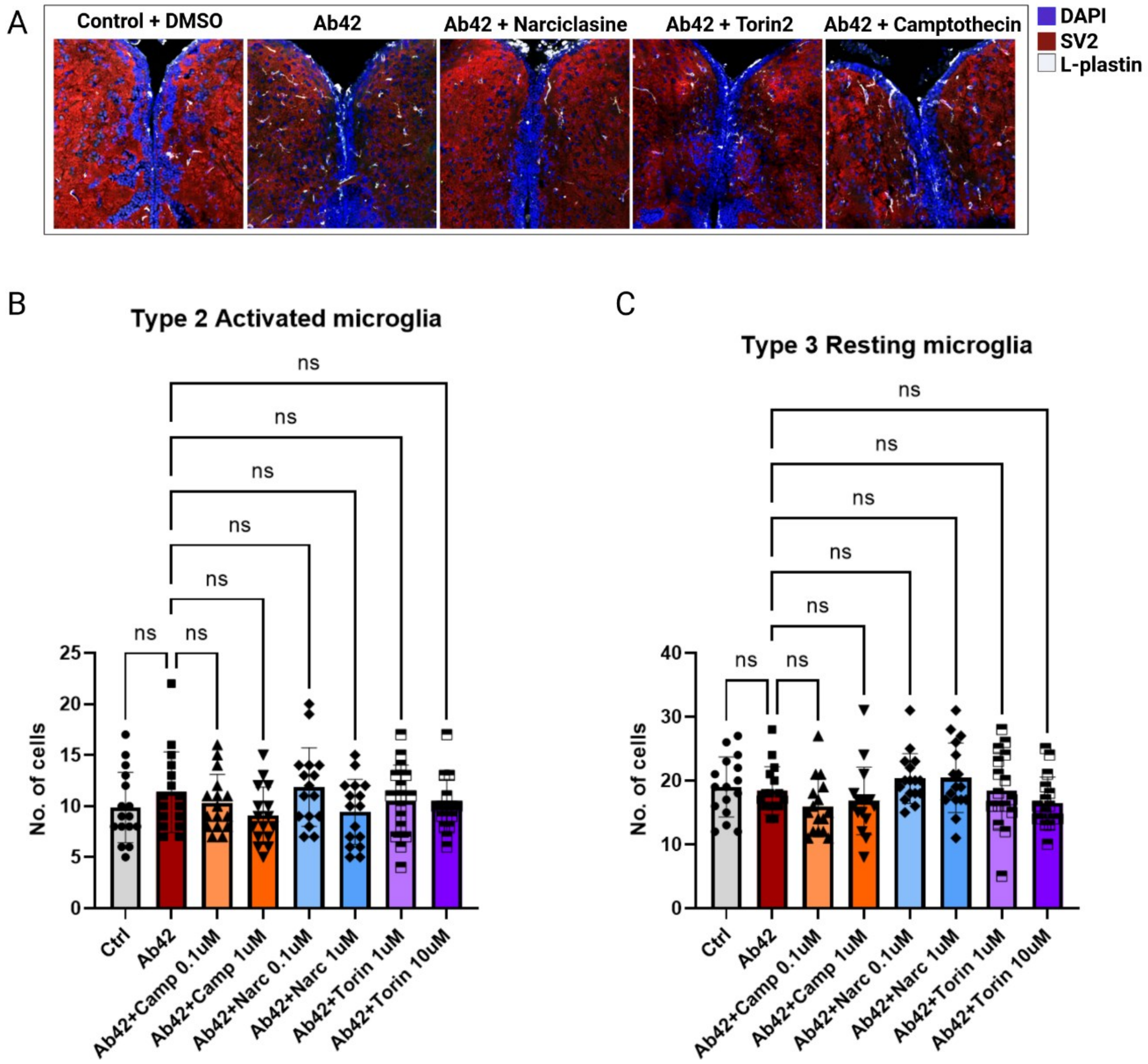


**Supplementary Figure 7. A.** Representative images of microglial- (L-plastin, white), synapse- (SV2, red) and nuclei (DAPI, blue) staining in the different treatment conditions Control + DMSO, Aβ42, Aβ42 + Narciclasine, Aβ42 + Torin2, Aβ42 + Camptothecin. **B-C.** Quantification of Type 2 (**B**) or Type 3 (**C**) activated microglia (L-plastin) was performed using confocal images of zebrafish brains harvested 5 days after Aβ42 injection plus compound- (Narciclasine: 0.1µM, 1µM; Torin2: 1µM, 10µM; Camptothecin: 0.1µM, 1µM) or DMSO-injection as control. Bar graphs represent mean cell number ± SD from a total of 16 images/ condition derived from 4 fish/condition. For statistical analysis, 2-way ANOVA followed by Dunnett's multiple comparison's test was performed. ns= non-significant.
